# Environmental microbiomes drive chemotactile sensation in octopus

**DOI:** 10.1101/2025.03.03.641191

**Authors:** Rebecka J. Sepela, Hao Jiang, Yern-Hyerk Shin, Jon Clardy, Ryan E. Hibbs, Nicholas W. Bellono

**Author notes:** These authors contributed equally to this work (Rebecka J. Sepela, Hao Jiang, Yern-Hyerk Shin).

## Abstract

Microbial communities coat nearly every surface in the environment and have co-existed with animals throughout evolution. Whether animals exploit omnipresent microbial cues to navigate their surroundings is not well understood. Octopuses use “taste by touch” chemotactile receptors (CRs) to explore the seafloor, but how they distinguish meaningful surfaces from the rocks and crevices they encounter is unknown. Here, we report that secreted signals from microbiomes of ecologically relevant surfaces activate CRs to guide octopus behavior. Distinct molecules isolated from specific bacterial strains located on prey or eggs bind single CRs in subtly different structural conformations to elicit distinct mechanisms of receptor activation, ion permeation and signal transduction, and maternal care and predation behavior. Thus, microbiomes on ecological surfaces act at the level of primary sensory receptors to inform behavior. Our study demonstrates that uncovering interkingdom interactions is essential to understanding how animal sensory systems evolved in a microbe rich world.

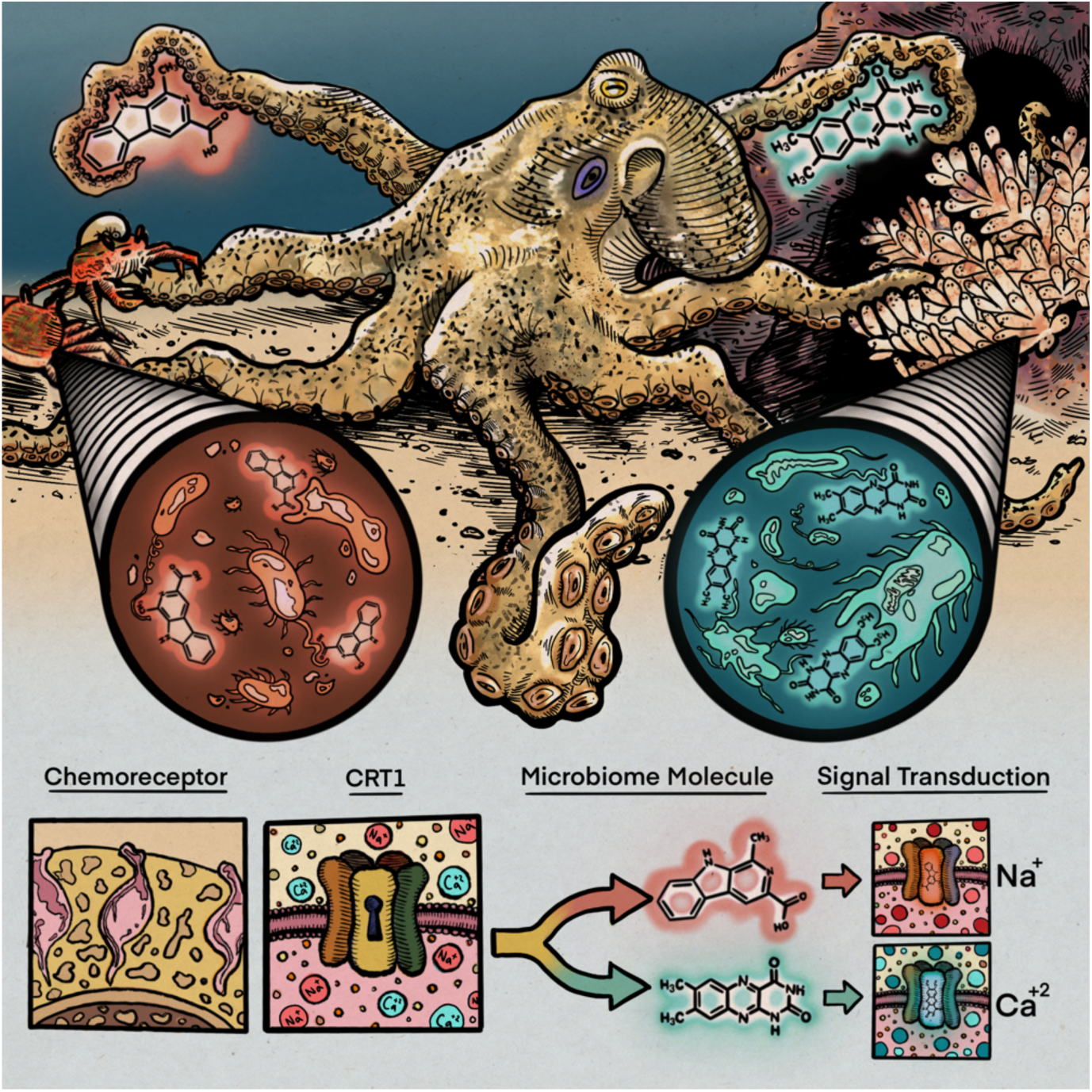

**Highlights:**

- Chemotactile receptors (CRs) detect microbiomes of prey and progeny
- Diverse microbial signals bind single CRs with distinct structural conformations
- Distinct microbial signals activate single CRs to permeate different ions
- Environmental microbes elicit octopus predatory and maternal behaviors

## Introduction

Three billion years after bacterial life originated, animals diverged from their protistan ancestors in seas teeming with bacteria but have remained in close association with these microbes throughout their history. The entirety of animal evolution has unfolded on a microbial stage, but we have only recently become aware of the effects of microbes on animal physiology^1,2^. Host-microbe symbioses span the tree of life and have afforded hosts the ability execute a broad repertoire of functions using microbial metabolic pathways^2^. The importance of host-microbe symbioses has now been demonstrated for numerous organ systems, including those for digestion, immunity, neural function, and the skin^3^. While the rudiments of host-microbe interactions are becoming clear, the process by which these pairings emerge is less well understood^4^. Just as microbes are motivated to find suitable hosts, there is increasing evidence that animals reciprocally deploy mechanisms that sculpt their interactions with microbes^2,4^. Yet the extent to which animals sense and respond to a rich microbial world and how they do so is not well understood^4^.

Microbes are well known for producing a vast array of diverse small molecules and are a defining feature of almost any organism or environment. Therefore, just as secreted microbial signals inform hosts of their internal microbiomes, microbe-derived chemicals could serve as a language for interkingdom signaling. Even the emergence of multicellularity in the closest ancestor to animals is triggered by microbial cues^3,5^, raising the intriguing possibility that microbial co-inhabitants may have evolved to influence complex animal behaviors including predation, predator avoidance, mating, and parental care^1,6^. Furthermore, microbiomes are influenced by environmental conditions such as changes in temperature, pH, sunlight, oxygen, and mineralized nutrient availability, positioning microbes as a fast-adapting, biotic interface to the abiotic world^7^. In this respect, the sensation of secreted microbial cues could reveal an otherwise imperceptible chemical world to diverse animals. Decoding this chemical language could reveal basic principles for how animals perceive their environment and how transient bacterial encounters transition to persistent symbioses^2^.

A major challenge in understanding chemosensation is identifying ecological ligand-receptor pairings in a world where chemical space is vast and diverse. Unlike vision or touch, the chemical world lacks clear organizational delineators and much of our knowledge has been restricted to large panels of synthetic compounds where the perceptive relevance is not always known. For this reason, studying the natural chemical constituents of the microbial world could provide important evolutionary insights into how animals evolved within their specific ecologies. This is especially true in aquatic environments, in which bacteria commonly accumulate at interfaces to form rich, surface-specific communities and produce surface-localized chemicals^9^. This raises the possibility that animals explore objects based on microbial chemical ecology, which could provide a natural chemical code for every surface they touch.

To test this hypothesis, we exploited octopus “taste by touch” sensation of seafloor surfaces as a model system. Octopuses use their flexible arms for contact-dependent chemotactile sensation of salient surfaces within cracks and crevices inaccessible to traditional sense organs^10^. Their unique body plan and large nervous system is organized such that most of the 500 million neurons are distributed along the arms to integrate peripheral signals and carry out autonomous search behaviors^11,12^. Octopus “taste by touch” is mediated by chemotactile receptors (CRs) that are enriched in the arm’s suction cups (suckers) and bind diverse ligands to transduce distinct electrical signals within the semi-autonomous arm nervous system^13^. CRs diverged from ancestral neurotransmitter receptors with novel structural adaptations to mediate contact-dependent chemosensation of surface-affixed, poorly soluble molecules that do not readily diffuse in water^14,15^. Yet, how octopuses distinguish biologically meaningful surfaces, such as their prey or eggs, from the crevices it explores is unknown.

What are the cues that define salient environmental stimuli? How is sensory receptor evolution shaped by microbiomes and their natural products? By profiling the culturable microbiome from octopus prey and progeny, we identify distinct crab and egg-derived microbial strains that produce specific molecules to bind octopus single CRs in different conformations. These molecules bias channel gating mechanisms such that diverse ligands trigger the same receptor to pass different ions and mediate distinct cellular signals. Environmental analyses show that these microbial cues are enriched in a surface-specific manner and we confirm they are sufficient to drive predation and maternal care behavior, demonstrating that animals use environmental microbiomes as important chemosensory cues.

## Results

### Microbiome signals activate octopus chemotactile receptors

To explore how octopuses distinguish objects in their environment, we first identified surfaces that elicit important behavioral responses. Octopuses are voracious hunters and opportunistic scavengers that use their arms to find hidden food when visual cues are limited^16^. In this case, they must distinguish live prey from decaying and potentially toxic food sources. Further, female octopuses exhibit acute awareness of their eggs by using their arms to clean and prune nonviable from viable embryos^17^. In both instances, octopuses use their arms to distinguish salient objects and then classify them as attractive or aversive via an unknown mechanism. Because octopus arms robustly respond to small molecule extracts from crabs and eggs (**Fig. 1A**, **Fig. S1A-D**), we analyzed their surfaces under different conditions that we suspected drive diverse octopus behaviors.

**Figure 1.**
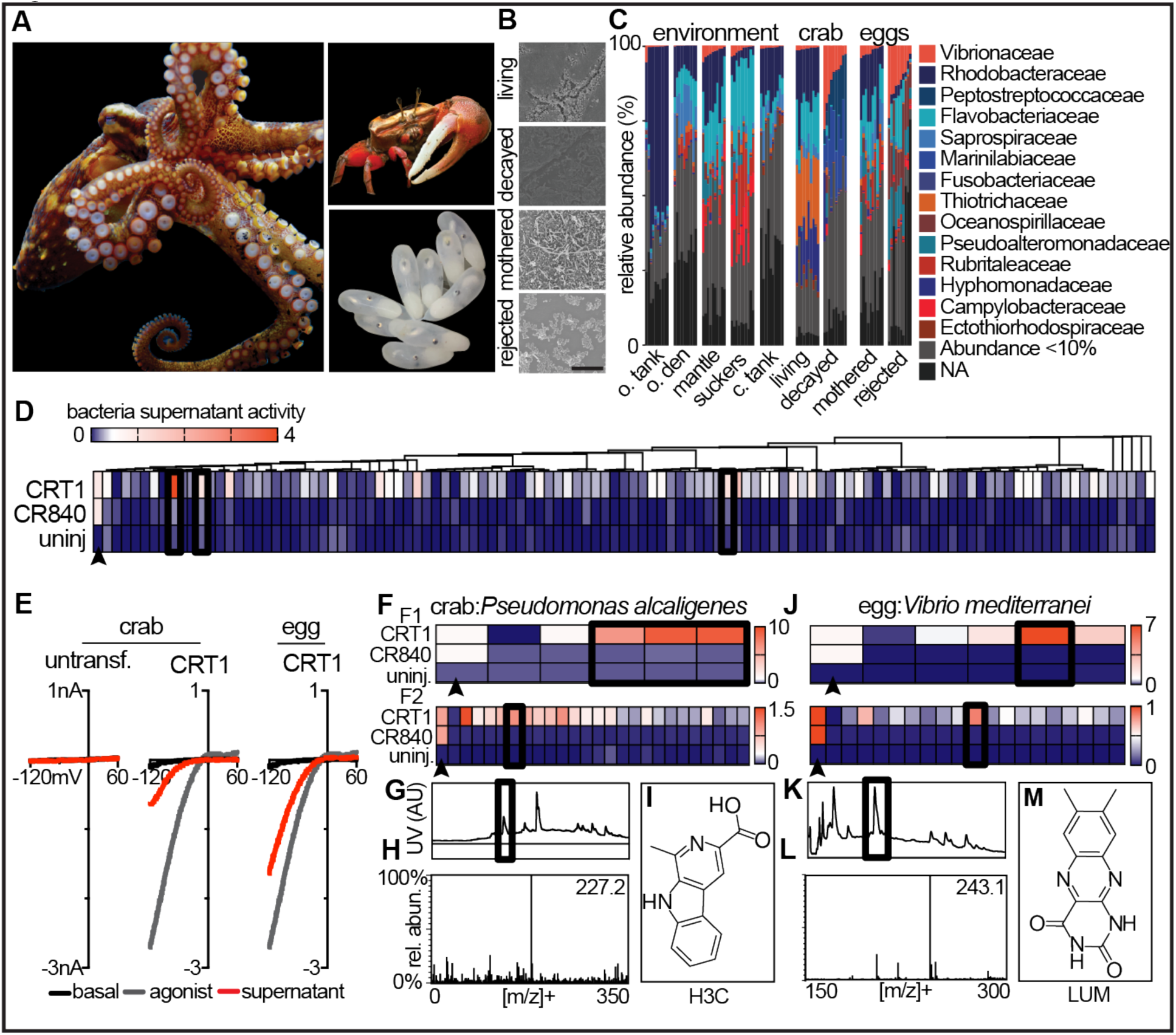
Environmental microbiomes produce chemical cues that activate octopus ‘taste-by-touch’ chemotactile receptors. **A,** *Octopus bimaculoides* displays its sensory suckers (*left*) next to salient cues including crab prey (*Leptuca pugilator, top right*) and developing octopus eggs (*bottom right*). **B,** Scanning electron micrographs reveal surface-adhered bacillus, coccus, and spirillum bacteria on crab shells and on octopus eggs; scale bar = 10 µm. **C,** Distinct microbiome communities were present on different surfaces from the octopus’s habitat, n = 10 for each surface. **D,** Culturable microbes were isolated from the carapaces of living and decaying crabs and mothered and rejected eggs. Supernatants from liquid cultures were filtered and screened for activity against a positive synthetic control agonist (black arrowhead) using TEVC from oocytes expressing octopus CRT1, CR840, and uninjected controls. Evolutionary relationships among strains were inferred by constructing maximum-likelihood phylogenetic trees. **E,** Select bioactive bacterial isolates (black boxes from *D*) from the oocyte screen were confirmed for CRT1 activity using patch-clamp recordings in HEK293 cells. **F & J**, Isolates cultured from crabs and eggs that showed activity in oocyte and HEK293 screens were iteratively sub-fractioned using HPLC and further screened using TEVC. **G & K,** UV chromatographs (360 nm) highlight the peak that corresponds with the most active subfraction. **H & L**, Mass spectrometry analysis reveals the major mass ions of the active fraction to be 227.2 and 243.1 [m/z^+^]. **L & M,** The major molecular components of the active fractions were harmane-3-carboxylic acid (H3C, from crab) and lumichrome (LUM, from egg), respectively.

Scanning electron microscopy revealed that crab shells and octopus egg casings harbor rich microbial communities that vary in morphology, abundance and composition based on condition (**Fig. 1B**). We observed that live crab carapaces were only sparsely populated with microbes, whereas those from decaying crabs were enveloped with coccoid-, rod-, filamentous-, and vibrio-shaped bacteria (**Fig. 1B**). Similarly, the microbial communities on mothered egg casings and rejected egg casings were distinct. The most pronounced differences were found in the morphology of the bacterial inhabitants, with rejected eggs housing spirillum-shaped bacteria. We hypothesized that differences in microbial populations, such as those found on live or decaying prey or mothered or rejected eggs, could influence how an octopus touches a surface. Consistent with this idea, octopuses differentially explored artificial surfaces affixed with cultured microbiomes from live and decayed crabs (**Fig. S1E-F**). 16s ribosomal rRNA V3-V4 barcoding confirmed that the taxonomic identities and abundances of the microbial communities of crabs and eggs were indeed state-dependent and also distinct from other objects in the octopus’s environment (**Fig. 1C, Fig. S1G,H, Table S1**). These results suggest that microbiomes constitute salient surfaces that could inform touch-dependent behaviors such as foraging or brooding.

We next asked if specific microbes from these ecologically relevant surfaces secrete molecules that are detected by the octopus chemotactile system. We isolated and cultivated ∼300 strains of marine bacteria from such surfaces to screen on cloned CRs for receptor activity. We used V1-V9 16S rRNA sequence barcoding to verify all strains as monocultures, collected supernatants, filtered for secreted small molecules (<3 kDa), and then tested for activity against octopus CRT1 and CR840 (**Fig. S1I-K**). CRT1 and CR840 represent the first cloned gene products from the 26-membered CR family. While CR subunits can combine to produce a diverse array of multimeric complexes, CRT1 and CR840 function as homomers and express well in heterologous systems^13,15^. Remarkably, this approach demonstrated that distinct strains activated CRT1 but elicited almost no activity with CR840 or uninjected oocytes as a control (**Fig. 1D**). Thus, distinct microbes activate specific sensory receptors.

Among tested microbial strains, CRT1 was robustly activated by *Pseudomonas alcaligenes* and *Vibrio alginolyticus* from crab shells and *Vibrio mediterranei* from incubated egg casings (**Fig. 1E** and **Fig. S1L**). CRT1 was previously shown to bind synthetic hydrophobic agonists, well suited to its role in contact-dependent chemosensation of relatively insoluble ligands by octopus arms^15^. Therefore, we hypothesized bacterial strains could provide a natural source for poorly soluble ligands and iteratively separated supernatants based on hydrophobicity to find that only select fractions activated CRT1. Active fractions were further sub-fractionated and analyzed with HPLC-MS and NMR to identify two β-carboline alkaloids, H3C (harmane-3-carboxylic acid) and 1A3 (1-acetyl-3-carboxy β-carboline) from bacteria isolated from crabs, and a flavin, LUM (lumichrome) from a strain harvested from eggs (**Fig. 1F-M, Fig. S1M-P, Fig. S2, Table S2**). Consistent with isolating these molecules from distinct microbes, they have different chemistries and can be produced downstream of distinct enzymatic pathways to carry out diverse functions including quorum sensing^18–22^. Indeed, we also found that these molecules differentially activated CRT1. Metabolites from crab microbes (H3C) had higher efficacy, but lower potency compared with those from egg (LUM) (**Fig. 2A, B, Table S3**). Neither H3C nor LUM produced significant channel desensitization. Importantly, both crab and egg-derived microbial signals elicited robust axial nerve cord activation and autonomous movement in isolated octopus arms, demonstrating that these molecules are detected by the octopus chemotactile system (**Fig. 2C, D**).

**Figure 2.**
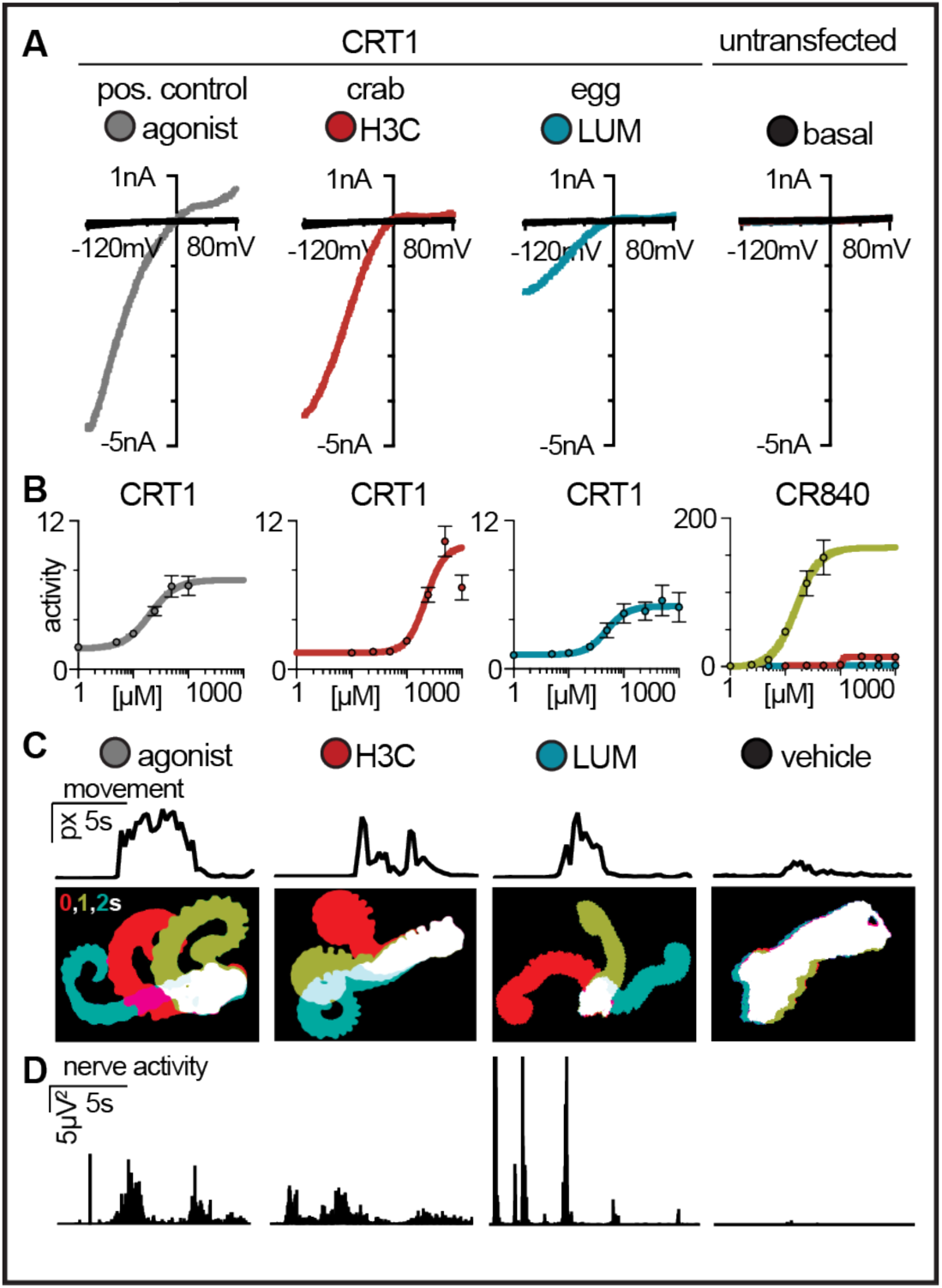
Microbial signals are pertinent chemical cues for the octopus sensory system. **A,** CRT1 responded to 500 µM H3C (crab, red), LUM (egg, blue), or a synthetic positive control agonist (nootkatone, grey). **B,** H3C was more efficacious while LUM was more potent. Neither chemical activated CR840, which was sensitive to a synthetic agonist, chloroquine (green). n = 3-7 cells per ligand. **C,** Octopus arms exhibit autonomous movement in responses to 1 mM H3C, 100 µM LUM, or 1 mM nootkatone but not vehicle control. *Top*: change in pixel values over time. *Bottom*: corresponding arm kymographs, red: 0s; green: 1s; teal: 2s. n = 8 arms. **D,** The axial nerve that innervates arm suckers was similarly activated by bacterial metabolites applied to suckers. n = 8-11 arms. See *Fig. S4* for quantitation of *C* & *D*. and *Table S3* for quantification of *B*.

### Structural basis of microbiome-driven sensation

We next exploited these single microbial-derived ligands to ask how sensory receptors evolve in the context of ecologically-relevant signals. Although LUM, H3C, and NOR (norharmane, a higher affinity structural analog of H3C) exhibited differences in efficacy and potency, cryogenic electron microscopy (cryo-EM) analyses revealed that all ligands were similarly bound to the hydrophobic orthosteric pocket. Furthermore, they all sampled numerous conformations, leading us to speculate that binding was mediated by nonspecific hydrophobic interactions (**Fig. 3A, Fig. S3A-I, Table S4**). The CR ligand binding pocket was previously found to diverge from ancestral nicotinic acetylcholine receptors to instead coordinate the poorly soluble synthetic ligands suited for contact-dependent aquatic chemosensation^15^. Indeed, the aromatic residues Y58, W122, F124, and F192 that coordinated all analyzed ligands were under positive evolutionary selective pressure and mutagenesis of these residues abolished ligand-evoked activity (**Fig. 3B, C, Fig. S3J, K**). To test whether ligand binding is purely mediated by hydrophobicity, we determined the apparent binding affinity of numerous structural analogs of identified microbial metabolites with varying hydrophobicity (**Table S3, Table S5**). Surprisingly, we found only a poor correlation between hydrophobicity and potency, suggesting that ligand recognition by CRs is more selective than could be explained by nonspecific hydrophobic interactions (**Fig. 3D**).

**Figure 3.**
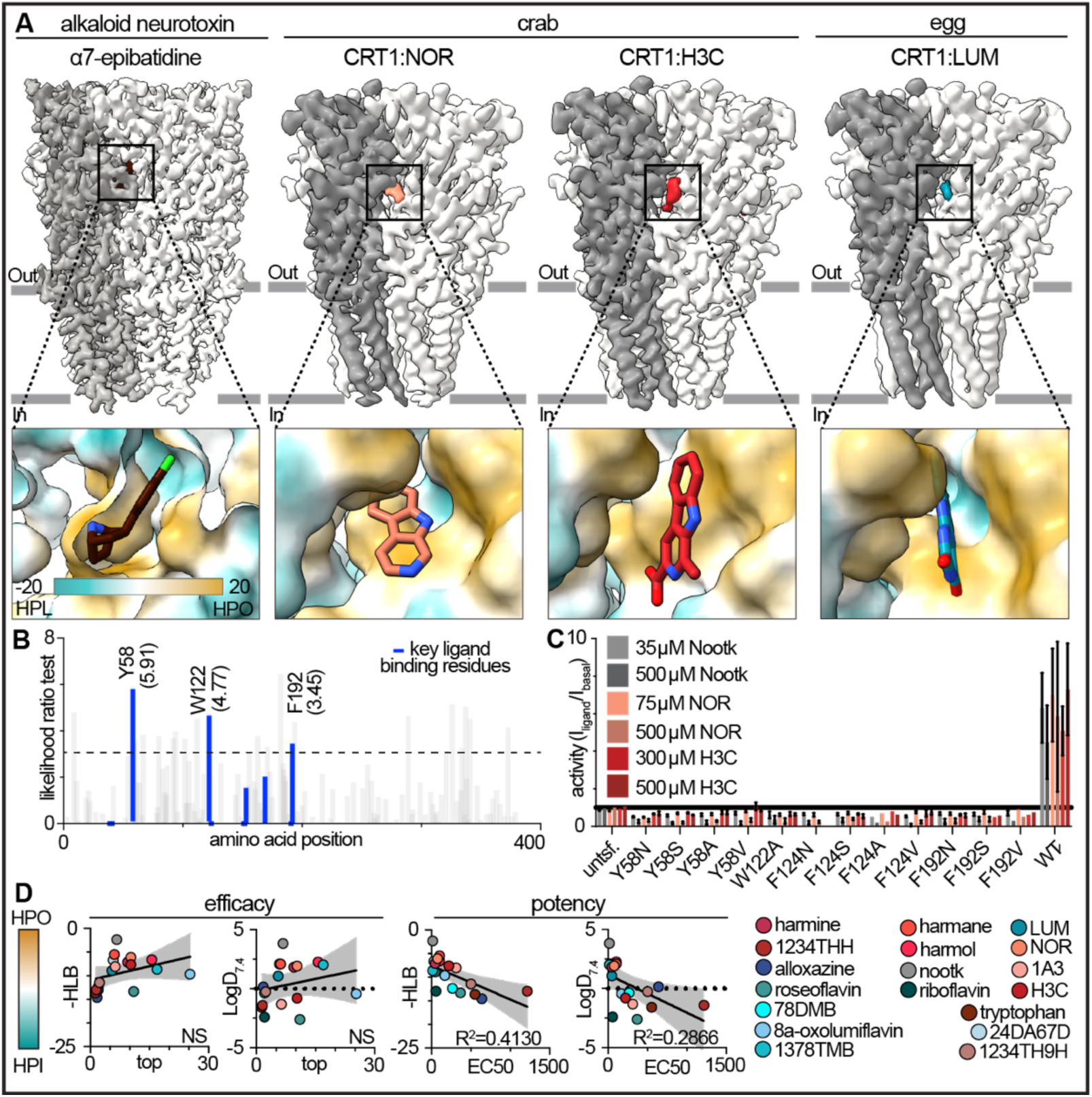
Octopus chemotactile receptors evolved a hydrophobic binding pocket to detect poorly soluble microbial metabolites. **A,** cryo-EM structures of CRT1 demonstrated that microbial ligands bind a flexible hydrophobic pocket (lower insets) in contrast to the tight coordination of ligands within the related human α7 neurotransmitter receptor binding pocket. All tested ecological ligands are positioned to form hydrophobic interactions within the CRT1 orthosteric binding site via residues Y58, W122, F124 and F192. **B,** Likelihood ratio test (LRT) for diversifying selection indicates that these residues are under positive selective pressure (blue, ≥ 3: p < 0.05). **C,** Site-directed mutagenesis of sites implicated in ligand binding resulted in a functional channel that was not activated by microbial ligands. n = 2-10. **D,** Despite the importance of hydrophobic interactions, neither the hydrophilic-lipophilic balance (HLB) nor the pH-adjusted water-octanol partitioning coefficient (LogD) of chemical analogs to H3C or LUM were strongly correlated with channel activation. This suggests that hydrophobicity alone does not mediate ligand binding. See *Table S3* and *Table S5* and *Figure S4* for channel activation metrics.

How do CRs differentially detect specific microbial signals to guide distinct octopus behaviors? In some chemosensory systems, different receptors detect structurally distinct chemical classes^23^. Yet here, ligands of distinct classes were bound in the same hydrophobic pocket of one receptor, intriguingly, with subtle nuances in binding poses. Microbial metabolites from crabs (H3C and NOR) were coordinated by a hydrogen-bond acceptor-donor interaction between the β-nitrogen of the β-carboline ring and the S169 side chain hydroxyl on CRT1 (**Fig. 4A, Fig. S3J).** This interaction was weaker in metabolites from egg (LUM, **Fig. 4A, Fig. S3J**). Could this hydrogen-bond contribute to differential sensation of microbiomes from crabs versus eggs? To ask this question, we performed another linear regression analysis of potency and efficacy versus a computational measure of the orbital electronegativity of the β-nitrogen of the crab microbiome-related molecules (**Fig. 4B, Table S3,** and **Table S5**). Here, we found a strong correlation between metabolite chemistry and receptor activity (**Fig. 4B-E, Fig. S4**). The most electronegative compounds also elicited octopus arm neural activity and autonomous movement (**Fig. 4B-E, Fig. S4**). Similar analyses with the egg microbiome-derived flavin molecules showed no such correlation, consistent with their binding lacking a strong hydrogen-bonding interaction (**Fig. S4**). To test whether this hydrogen-bond was important in specifying binding of crab-versus egg-derived microbial signals, we first abrogated the hydrogen-bond donor capabilities of CRT1 by mutagenizing S169 into alanine. This mutation markedly reduced potency for NOR (crab) but did not significantly affect the apparent affinity of LUM (egg, **Fig. 4F**). We then tested the serine hydrogen donor on the crab microbial metabolites by comparing the binding affinity of NOR with carbazole, a NOR analog that lacks a β-nitrogen and thus cannot accept a hydrogen bond (**Fig. 4G**). Unlike NOR, carbazole was unable to elicit receptor activity, demonstrating the importance of specific receptor-ligand interactions for microbiome signals (**Fig. 4G**).

**Figure 4.**
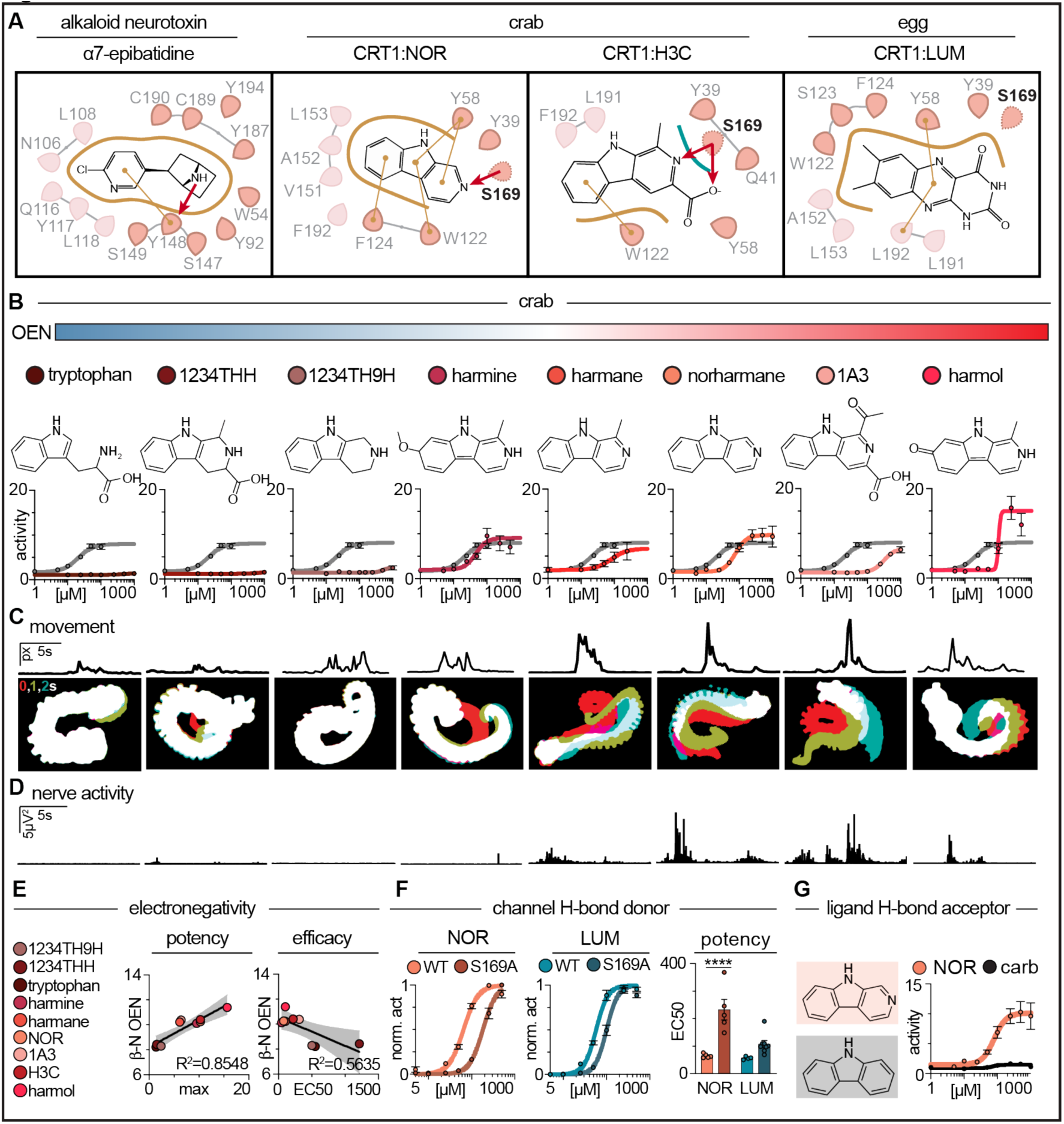
Crab- and egg-derived microbial metabolites exhibit distinct receptor binding mechanisms. **A,** Crab (H3C and NOR) and egg-derived (LUM) microbial metabolites relied heavily on hydrophobic interactions (thick gold lines) for CRT1 coordination; thin gold lines indicate aromatic-π stacking. Only crab-derived microbial ligands additionally engaged in H-bonding interactions at S169 (dark red arrows). **B - D,** Analogs to crab-derived microbial signals with increasing orbital electronegativity (OEN) evoked strong activity in expressed CRT1, autonomous octopus arm movement, and axial nerve cord activity in response to stimulated suckers. n = 3-7 cells per ligand, n = 3-8 arms. **E,** Crab-derived microbial ligand potency and efficacy were tightly correlated with the OEN of the ligand’s tertiary nitrogen (potency: p = 0.0198, efficacy: p = 0.0004). **F,** Mutation of H-bond donating S169 reduced apparent affinity of NOR but not LUM (n = 4-8, p < 0.0001, one-way ANOVA with Sidak’s multiple comparisons test). **G,** Replacement of the H-bond accepting β-nitrogen of NOR with a carbon atom preserves molecular hydrophobicity but still abolishes CRT1 binding capacity, demonstrating the essential H-bond interaction (n = 4-5, p < 0.0001, one-way ANOVA with Sidak’s single pooled variance comparisons test). See *Table S3* for pharmacological fit values; *Fig. S4* for arm behavior and nerve quantitation. See *Table S5* for chemical metrics.

Activation of CRT1 leads to direct ion permeation through its central pore to elicit downstream signaling events, such as the initiation of neural activity. The ion permeation pathway includes an extracellular vestibule lined with charged residues that tune conductance. We previously found that CRT1 uses E104 to form a negatively charged ring in the extracellular vestibule that facilitates high permeation of calcium, an essential second messenger^15^. Curiously, crab- and egg-derived microbial ligands stabilized distinct structural conformations of the Ω-loop that contains E104. With egg-derived LUM bound, E104 oriented toward the central channel axis, ideally positioned to bias calcium permeation through the open pore **(Fig. 5A**, **Fig. S5A**). However, when CRT1 was bound to crab microbial metabolites NOR or H3C, E104 retracted away from the open pore (**Fig. 5B, Fig. S5B,C**). Since the map density of E104 and its Ω-loop suggested a mixture of conformations (**Fig. S5D-F**), we tested the relative calcium permeability in the presence of these distinct agonists. Consistent with structural observations, we found the egg-derived agonist LUM elicited higher calcium permeation relative to the crab-derived molecule NOR (**Fig. 5C,D**). Furthermore, mutating E104 to alanine reduced calcium permeability evoked by LUM, but did not affect NOR (**Fig 5D**). Thus, ligands from microbiomes of different ecologically relevant surfaces mediate distinct cellular signals through the same receptor through stabilizing alternative conformational states. This local mechanism of signal coding at the level of the primary sensory receptor is well suited to contribute to peripheral processing in the distributed, semi-autonomous octopus nervous system.

**Figure 5.**
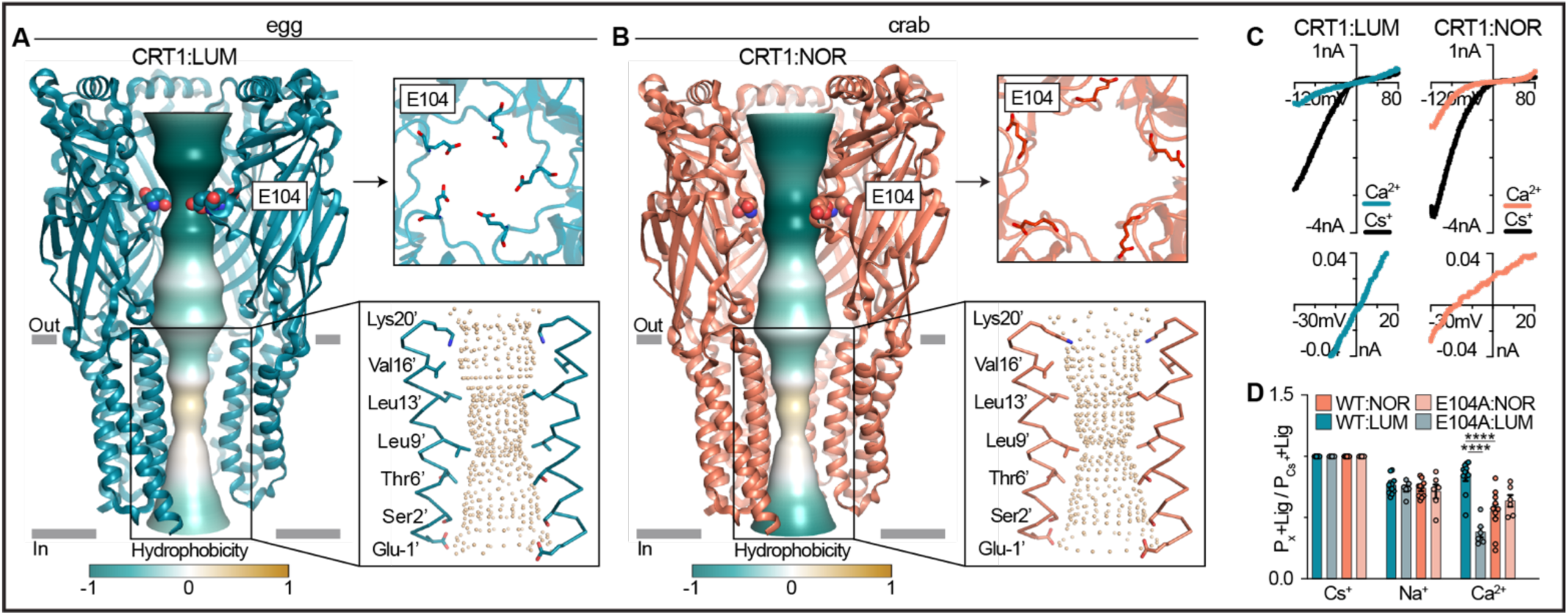
Microbial metabolites from crabs and eggs differentially tune chemotactile receptor ion permeation. **A,** Ion permeation pathway of CRT1-LUM complex (colored by hydrophobicity). Top-down view illustrates that residue E104 pointed toward the center of pore to form a constriction in the extracellular portion of the permeation pathway. *Inset:* The pore-lining residues of two M2 helices are shown as sticks, with spheres indicating an open pore. **B,** The cytosolic side of the ion permeation pathway of the CRT1-NOR complex was similar to CRT1-LUM, but E104 rotated away from the pore axis. *Inset:* The CRT1-NOR complex similarly exhibited an open pore. **C,** LUM and NOR activated distinct channel permeation indicated by shifted reversal potentials observed in current-voltage relationships. **D,** LUM evoked significantly higher Ca^2+^ permeability than NOR that was reduced by mutating E104. Cs^+^ and Na^+^ permeation were similar and NOR Ca^2+^ permeability was unaffected by E104A. (n = 6-11, p < 0.0001, one-way ANOVA with Sidak’s multiple comparisons test).

### Microbial chemical ecology drives surface-specific sensory behaviors

We next sought to understand how microbiome-derived ligands elicit distinct cellular signals to drive organismal behavior. We first revisited our environmental microbiome analyses to determine the abundance of bioactive strains on different surfaces, thereby indicating where microbial ligands could be sensed by exploring octopuses. Although microbe strains of interest were initially associated with crab and egg surfaces, culturing-based approximations of a microbial community are not sufficient to capture the relative abundance of a strain within and between specific microbiomes^24^. Out of 12,343 total amplicon sequencing variants (ASVs) that represent distinct bacterial strain clusters from the amalgamation of ten different surface-specific microbiomes, 52 ASVs shared >99% sequence identity to the cluster of *Vibrio alginolyticus* strains (isolated from crab) and 104 ASVs matched the *Vibrio mediterranei* cluster (from egg). *Vibrio alginolyticus* ASVs accounted for at least 7% of the total bacterial community on the decaying crab carapace, which changed over the course of decay (**Fig. 6A**, **Fig. S6A-D**). Similarly, *Vibrio mediterranei* was overrepresented in the microbial community growing on rejected eggs. Further, *Vibrio alginolyticus* and *Vibrio mediterranei* were among the most differentially prevalent strains for the decayed crab and rejected egg surfaces, respectively (**Fig. 6A**, **Fig. S6E**). From these microbiome analyses, we suspected that the molecules produced by these microbes are enriched in decaying crab shells and rejected octopus eggs, respectively. Indeed, increased fluorescence of these surfaces correlated with the inherent fluorescence of H3C and LUM (**Fig. 6B**). Importantly, we also directly detected H3C and LUM from crab carapaces and egg casings at concentrations sufficient to evoke CRT1 activity (**Fig. 6C**). Intriguingly, 53% of decaying crabs tested possessed detectable H3C, compared to only 20% of living crabs. LUM was detected on both crab and egg surfaces but was three to four times more abundant in decaying crabs and rejected eggs than their respective counterparts (**Fig. S6F-K**). These results demonstrate that microbiomes and their bioactive molecules are present in specific ecologically relevant surfaces important for behaving octopuses.

**Figure 6.**
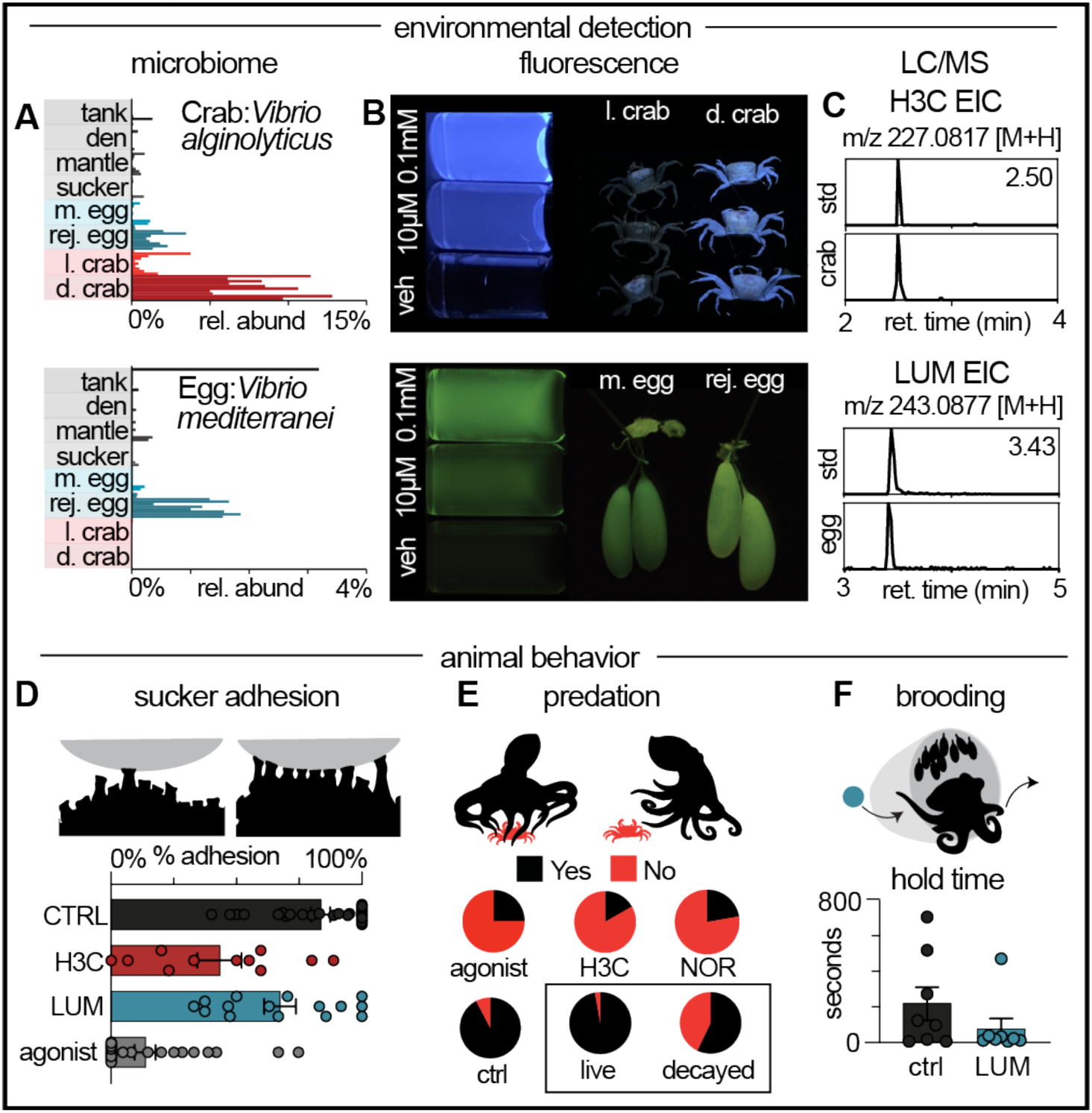
Microbial metabolites are present in the octopus’ environment and drive distinct chemotactile behaviors. **A,** H3C-producing *Vibrio alginolyticus* was enriched in the decaying crab microbial community and LUM-producing *Vibrio mediterranei* was enriched in rejected egg communities, respectively. **B,** The surface of decayed crabs and eggs fluoresced when illuminated by long-wave UV-light, similar to that produced by pure H3C and LUM. **C,** H3C and LUM were detected from environmental samples as evident by a matched LC/MS elution time of the major [M+H]^+^ ion (bottom) with a commercially available standard (top). **D,** Autonomous arm behavior showed that sucker adhesion was decreased by coated surfaces with 1 mM H3C (p = 0.0016), 500 µM LUMI (p = 0.0330), or 1 mM nootkatone (p < 0.0001) compared with DMSO as a control (n = 11-15 arms, Geisser-Greenhouse correction and Dunnett’s multiple comparisons test). **E**, Hungry octopuses investigated all crab mimics and engulfed those treated with 10% DMSO (n = 13 trials), similar to real immobilized crabs (n = 35). Octopuses did not engulf mimics treated with 1 mM H3C (n = 12) or 10 mM nootkatone (positive control agonist, n = 8), similar to how octopuses reject real decayed crabs (n = 21). **F**, Brooding octopus moms rejected egg mimics containing 500 µM LUM from their clutch faster than vehicle mimics (1% DMSO, n = 8, p = 0.0391, Wilcoxon matched-pairs signed rank test).

Considering that the bioactive crab-derived microbe and metabolite H3C was enriched in a decaying state and that the egg-derived microbe and metabolite LUM was seen in higher abundance in rejected eggs, we wondered if these signals might activate CRT1 to encode and elicit an aversive response. Even when removed from the octopus, suckers retain the ability to adhere to surfaces, exerting stereotyped degrees of force that reflect the surface presented^25^. To test if microbial signals affect sucker adhesion, we presented isolated arms with agar-coated coverslips doped with seawater, H3C (from decaying crab microbes), LUM (from rejected egg microbes), or the synthetic CRT1 agonist nootkatone. Consistent with aversive autonomous sucker behavior, H3C and nootkatone significantly reduced adhesion compared to control, while the effect of LUM was similar but more subtle (**Fig. 6D, Fig. S6L**). These results show that microbial signals drive reflexive aversive behavior, even at the level of single suckers.

How are distinct microbiome signals encoded to elicit specific chemotactile behaviors? To ask if H3C is sufficient to drive differences in predatory behavior of live versus decaying crabs, we soaked crab mimics in H3C, NOR, or nootkatone and vehicle controls and compared octopus behavioral responses to those elicited by real crabs. Animals investigated mimic crabs the same as real immobilized or decaying crabs, likely based on visual cues (**Fig. S6M**). However, following first touch, octopuses only readily engulfed vehicle-treated mimic crabs the same as real crabs (**Fig. 6E, Video S1**). In contrast, following first touch, they avoided those soaked with the microbial signals H3C, NOR, or the CRT1 agonist nootkatone, similar to how they responded to decaying crabs (**Fig 6E, Fig. S6M, Video S2**). In parallel, we asked whether the microbial signal LUM was sufficient to induce egg rejection behavior by brooding octopuses. For this assay, we presented DMSO and LUM-treated egg mimics to octopus moms actively brooding their clutch. Following presentation of vehicle or LUM-doped egg mimics into their den, octopuses groomed the mimics similar to how they treat their own eggs and then eventually discarded them (**Video S3**)^17^. In this case, brooding octopuses ejected LUM eggs significantly faster than vehicle eggs, consistent with this signal being sufficient to induce egg rejection (**Fig. 6F, Video S4**). Thus, microbial signals are sufficient to drive specific chemotactile behaviors.

## Discussion

Every organism experiences the same world differently based on their ability to extract cues that are most salient to their biological needs^26^. Therefore, understanding how organisms evolve to occupy distinct niches requires discerning the natural signals produced in their environment. While chemical space has historically been defined by the number of synthesizable or extractable molecules, ancestral chemical space was forged by microbes. The octopus represents a powerful model to probe this hypothesis because the octopus’s chemical world is defined by surface-affixed compounds that direct CRs to transmit signals through their complex nervous system. Octopus arms contain the majority of their neurons, allowing for local signal detection, processing, and behavior that facilitates prey capture along the seafloor^27^. Here, we report the discovery of molecules from discrete microbiomes that octopuses detect among complex natural environments. By exploiting these ecologically relevant cues, we show that a single sensory receptor possesses structural adaptations to differentially sense and transmit signals that indicate distinct behaviorally meaningful cues. Not only do prey and egg-derived molecules bind in different conformations to modulate potency and efficacy, but they also result in distinct channel gating mechanisms such that different ligands trigger the same receptor to pass different ions. Thus, even the primary sensory receptor encodes complex information about the environment by eliciting specific cellular messages, well suited to the distributed nervous system of the octopus.

Prior to this study, the valence encoded by the octopus chemotactile system was unknown. Here, we discover that microbial ligands encode aversive cues for both predation and brooding. Octopuses are opportunistic foragers that use “blind feeding” exploration among seafloor crevices^10^. These opportunistic hunters predominantly forage in crevices and at night, where visual cues are limited^10^ and microbial cues that act on the chemotactile arms could provide a “taste by touch” signal to deter animals from feeding on decaying and potentially toxic prey. Indeed, we find that microbial CR agonists are enriched in decaying crabs, which could help inform octopuses to abort search behavior. Octopuses also play an instrumental role in developing their eggs. Many species cement their eggs onto hard substrates within a den, guard and care for the clutch by continuously cleaning and aerating the eggs with their suckers^17,28^. Even the non-brooding females will carry out egg grooming behavior when presented with a clutch from another mother, suggesting eggs are potent sensory cues^17^. Egg-brooding is commonly the final act of a female octopus’s life, yet these females will remove select eggs from the clutch as infertile or dead embryos, suggesting a “taste by touch” mechanism for distinguishing egg health in maternal care^17^. In agreement with this hypothesis, we find that microbial cues are enriched on rejected eggs. While these two examples demonstrate how microbes produce ecologically relevant ligands to inform sensory behavior, microbiomes exhibit complex chemical interplay and time-dependent dynamics that can affect their metabolomes. Therefore, a myriad of unknown natural compounds, receptors, and behavioral contexts involving microbe-animal interactions await description.

Chemosensation is an ancient sensory modality that evolved within a landscape saturated by microbial communities. Indeed, chemosensation was an essential evolutionary precursor to multicellular organization and intracellular communication, and likely sculpted the rules of engagement for cross-kingdom interactions^29^. Our work suggests that footprints of this intertwined past not only define symbiosis and internal host physiology, but also the sensory experience of animals. Indeed, this microbial basis of sensory biology likely spans the tree of life. As such, examples can be found in fly foraging and oviposition^30^, marine invertebrate larvae settlement^31^, animal predation of bacteria^32^, and mammalian mate selection^33^. Discovering and comparing diverse examples of additional cross-kingdom signals will be essential to understanding the evolutionary transition of environmental sensation of the microbial world to the formation of long-lasting symbioses. Ultimately such approaches will illuminate principles regarding the molecular language of interkingdom symbiosis and co-evolution.

## Acknowledgements

We thank C. Winkler for providing octopuses; B. Walsh, P. Kilian, A. Moulton, N. Mongillo, T. Hautala, M. Saltzmann, C. Ammann, J. Green, J. Thibodeau for assistance with animal care and behavioral experiments; F. Muro Villanueva and R. Nett for assistance with mass spectrometry measurements; Lily Soucy and Anik Grearson for illustrations and photos; P. Vaelli for contributions to the early stages of the project; the HMS Electron Microscopy Facility for imaging and consultation services; This research was further supported by grants to N.W.B. from the New York Stem Cell Foundation and NIH (nos. R35GM142697 and R01NS129060), NIH grants to R.E.H. (R01NS129060) and J.C. (R01AI172147), and an NRSA fellowship to R.J.S (F32GM148163).

## Author contributions

Conceptualization: RJS, NWB; Data curation: RJS, YHS, HJ; Formal analysis: RJS, YHS, HJ; Funding acquisition: NWB, JC, REH; Investigation: RJS, YHS, HJ; Methodology: RJS, YHS, HJ; Project administration: RJS, NWB; Software: RJS, YHS, HJ; Resources: RJS, YHS, HJ; Supervision: NWB, REH, JC; Validation: RJS, YHS, HJ; Visualization: RJS, YHS, HJ; Writing – original draft: RJS, NWB; Writing – review and editing: all authors.

## Declaration of generative AI and AI-assisted technologies

Generative AI and AI-assisted technologies were not used in the preparation or production of this manuscript.

## Supplemental Information titles and legends

Document S1. Figures S1-S6. Tables S1-S6.

Video S1. Octopuses will engulf crab mimics soaked in DMSO vehicle, related to *Figure 6E*.

Video S2. Octopuses do not engulf crab mimics soaked in crab-like microbial metabolites, related to *Figure 6E*.

Video S3. Octopus moms quickly reject agar eggs treated with egg-derived microbial metabolites (LUM), related to *Figure 6F*.

Video S4. Octopus moms tolerate agar eggs prepared with the vehicle DMSO, related to *Figure 6F*.

## STAR methods

### Key resources table

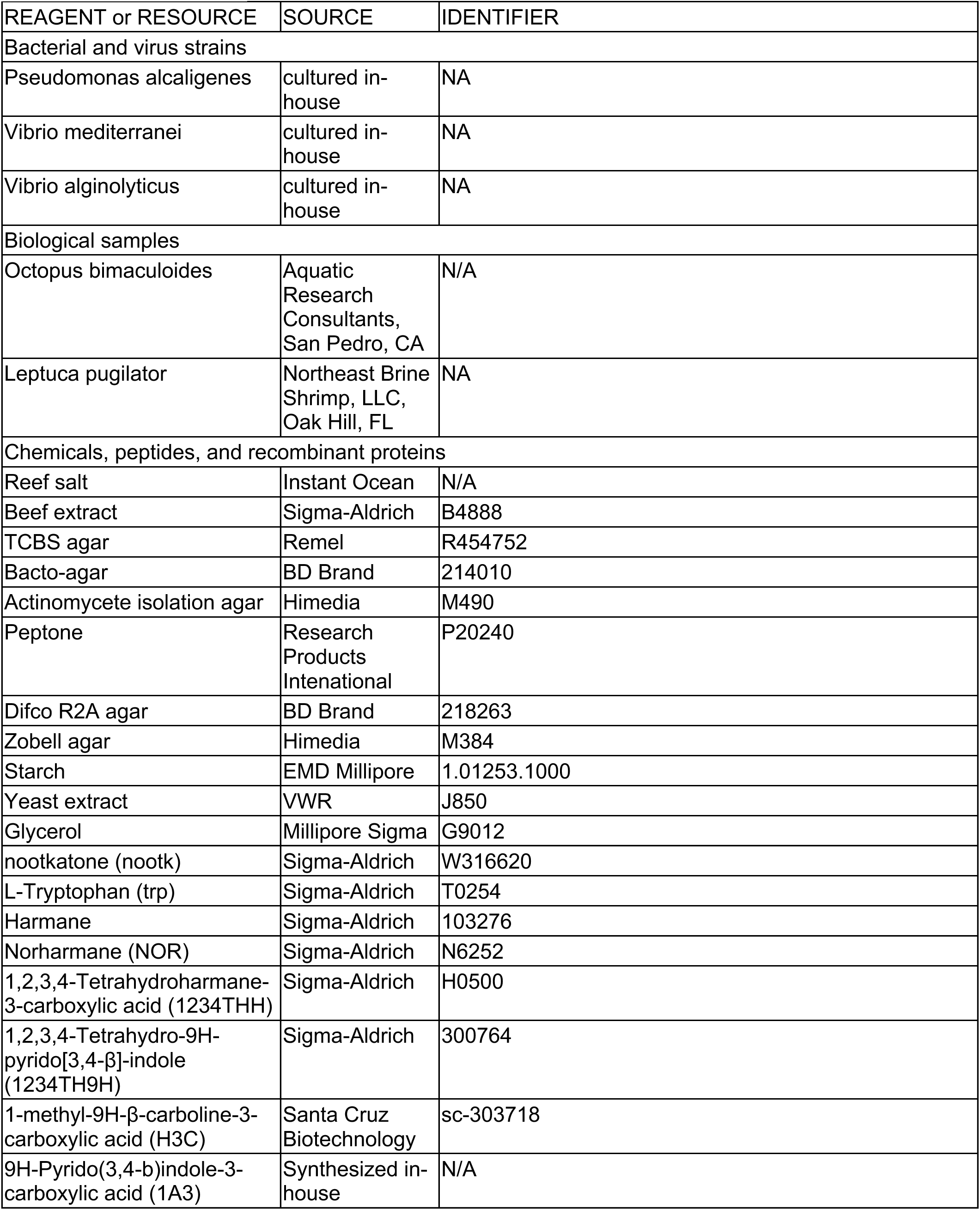
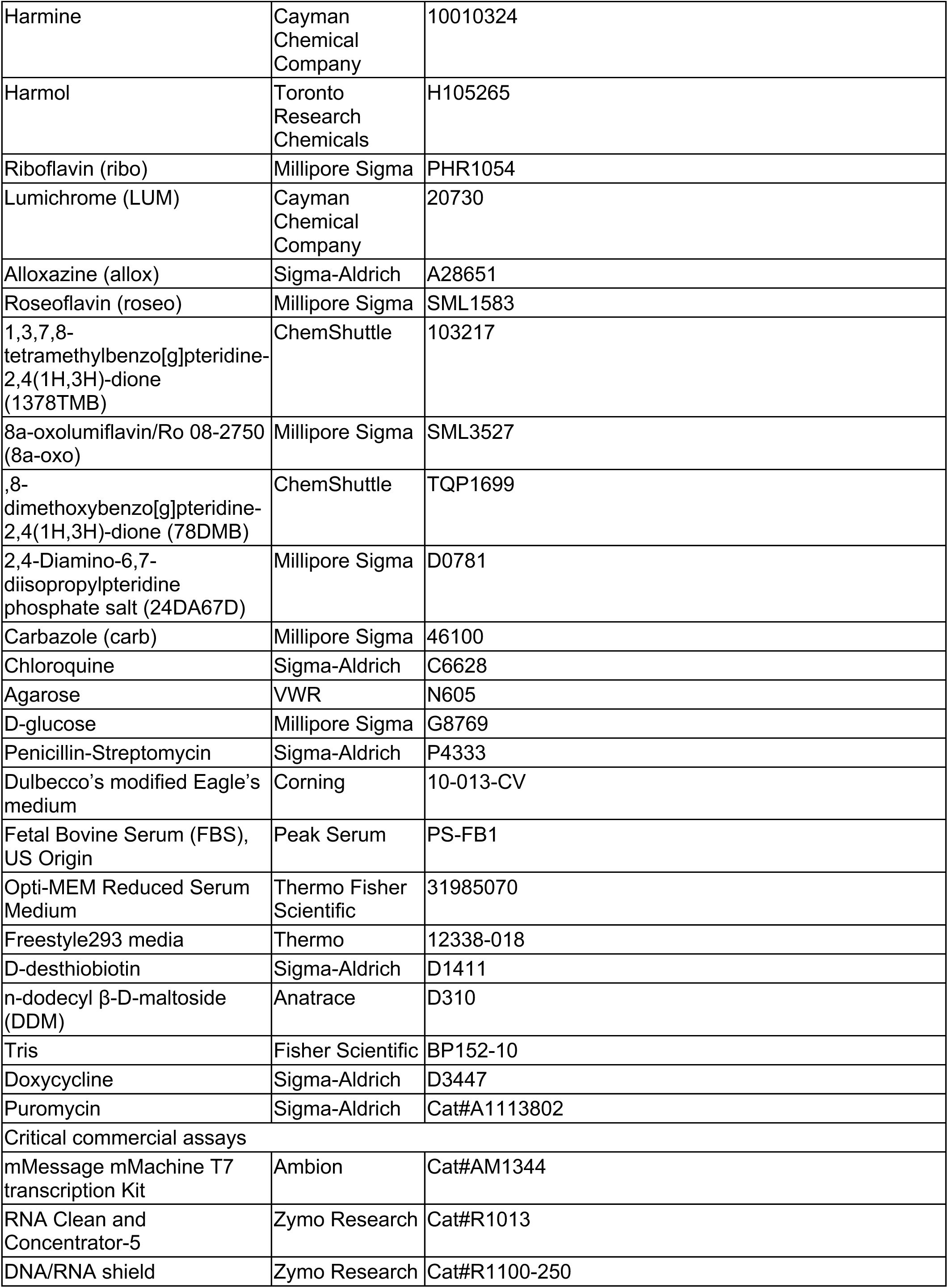
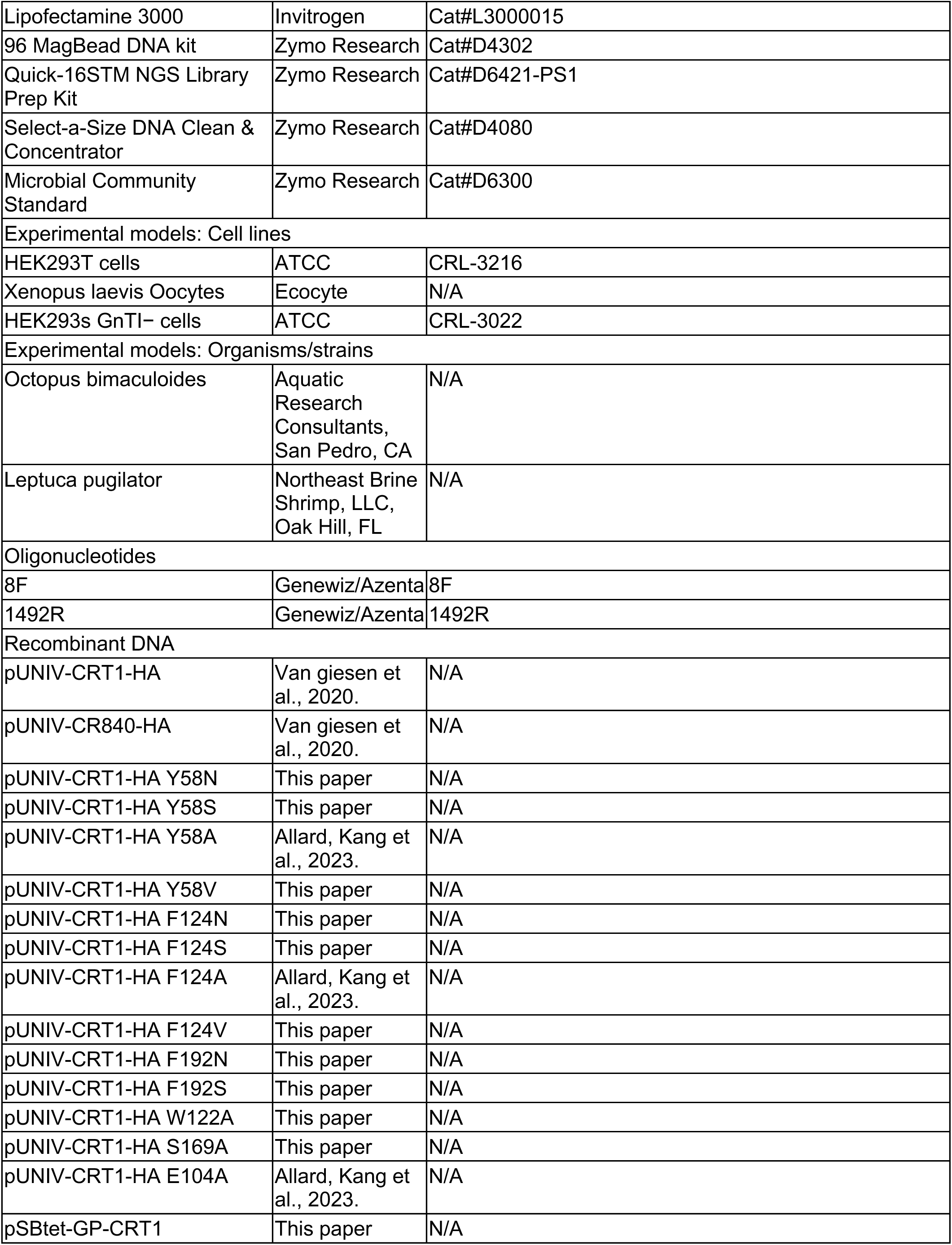
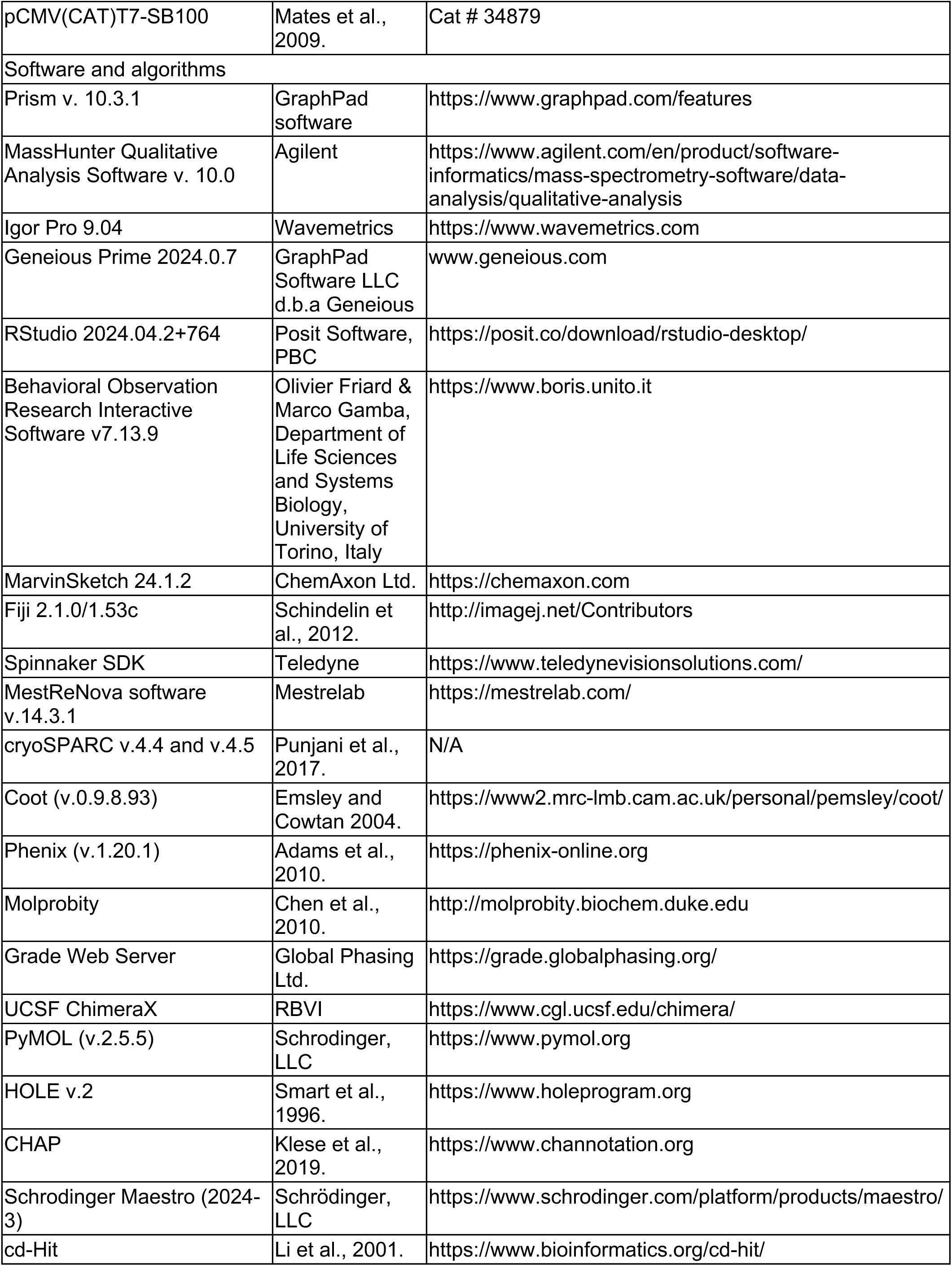
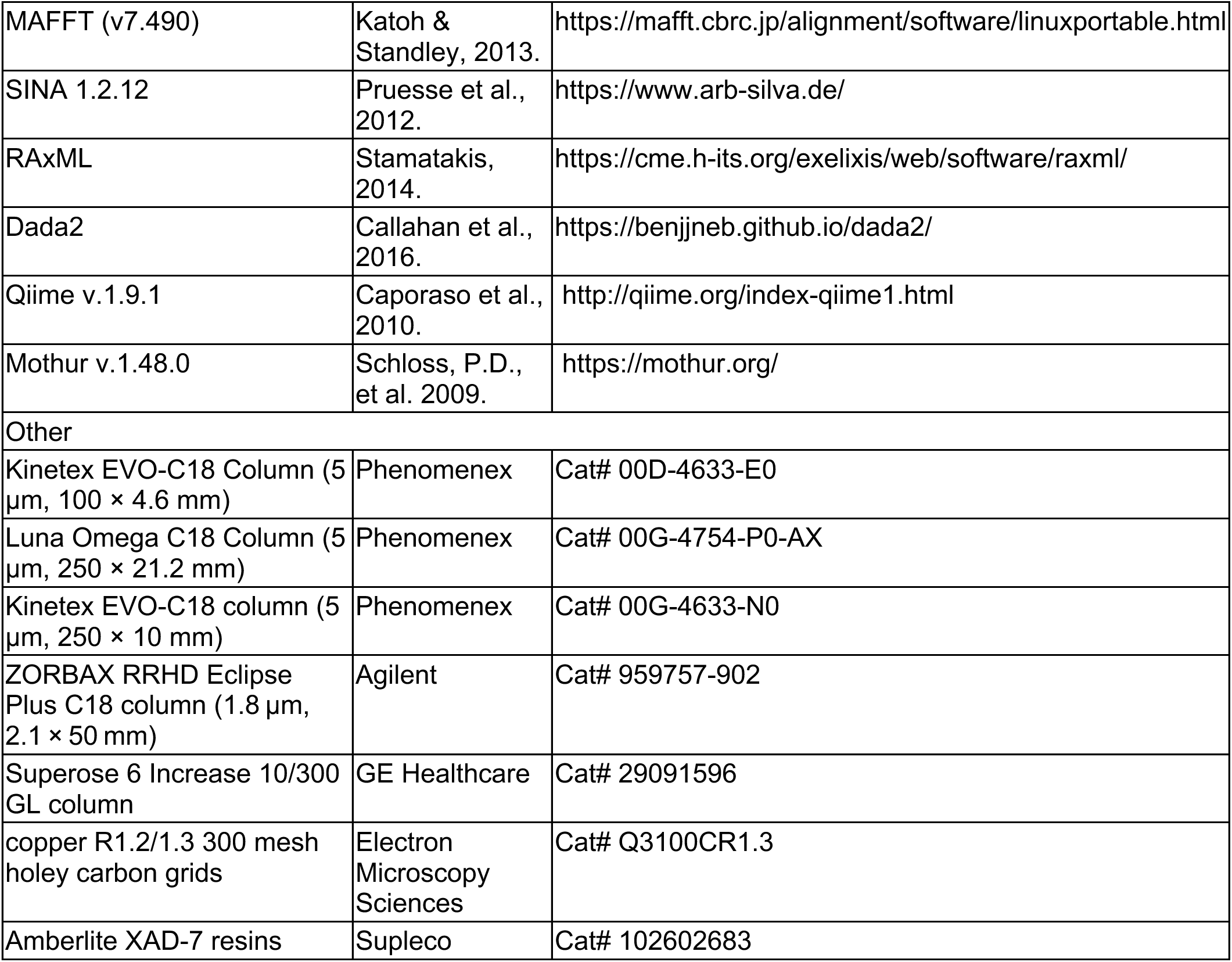

## Resource availability

### Lead contact

Further requests for resources and reagents should be directed to and will be fulfilled by the lead contact, Nicholas Bellono.

### Materials availability

All unique reagents generated in this study are available from the Lead Contact.

### Data and code availability

Atomic model coordinates and cryo-EM density maps for the CRT1 structures bound to NOR, H3C or LUM have been deposited in the Protein Data Bank with accession code 9E6B, 9E6C, 9E6D and in the Electron Microscopy Data Bank with accession code EMD-47554, EMD-47555, EMD-47556, respectively.

## Experimental model and subject details

California two-spot octopuses (*O.bimaculoides*) were wild-caught male and female adults (1-2 years old, Aquatic Research Consultants). Expect where noted, animals were fed daily with fiddler crabs (*Leptuca pugilator,* Northeast Brine Shrimp) and kept on a 12-h light-dark cycle in natural seawater. Animal protocols were approved by the Harvard University Animal Care and Use Committee (protocol ID: 18-05-325).

### Method details

#### Microbiology

##### Microbiome scanning electron microscopy

Scanning electron micrographs of *Leptuca pugilator* shells (fixed immediately and 24 hrs following pithing) and *Octopus bimaculoides* eggs (harvested from brooding mothers or retrieved from the tank a maximum of 24 hours after being expelled from the clutch) were acquired by the Harvard Medical School Electron Microscopy Core. Briefly, after being fixed in formaldehyde/glutaraldehyde 2.5% in 0.1M sodium cacodylate (Electron microscopy services, Cat#15949) for >2 days. Samples were rinsed three times, 10-15 min each time, in 0.1M sodium cacodylate buffer, pH 7.4, then soaked for 1-2 hrs in 1% osmium tetroxide in 0.1M sodium cacodylate buffe, pH 7.4. Samples were rinsed three times, five minutes each, in water and then dehydrated in ethanol using the following soaking progression: 50% EtOH, 5min; 70% EtOH, 5min; 95% EtOH, 10 min; 100% EtOH, 10 min; 100% EtOH, 10 min. Samples were then dried by Critical Point Dry (45-60min) and mounted onto stubs using conductive carbon tape, silver paint or paste and then sputter coated at a thickness of 1-5 nm. Images were taken using S4700 at 5.0kV 10.7 mm x 2.00k.

##### Microbiome sampling and isolate cultivation

Bacterial samples were collected using sterile, individually packaged, cotton-tipped applicators (McKesson, MFR #24-106-1s). Collection sites and subjects were first rinsed in reverse osmosis H_2_O to remove transient bacteria and then swabbed for ∼10 s using a rotating motion. Rejected eggs were swabbed a maximum of 12-24 hours following ejection. Unincubated eggs were removed from an actively mothered clutch and allowed to remain in the tank for 5 days before swabbing. Live fiddler crabs were sampled from the top and bottom carapace for 5 seconds on each side. Decaying crabs were sampled following a 1-8 day period of room temperature decay in stagnant, 0.2 µm-filtered seawater. For V3-V4 microbiome sequencing, swabs were placed in cryovials containing 100 µL DNA/RNA Shield^TM^ (Zymo Research, Irvine, CA) and stored at -80 °C. For preservation of community microbiomes, swabs were placed in cryovials containing equal parts seawater and 2x freezing stock solution (20% DMSO, 30% seawater, 50% glycerol). For isolate cultivation, swabs were soaked in 1 mL seawater for 5 minutes and diluted ten-fold over four serial dilutions (10^0^, 10^-1^, 10^-2^, and 10^-3^). 100 μL of each dilution was plated onto TCBS agar (Remel, R454752; prepared by manufacturers’ protocol), actinomycete isolation agar (Himedia, M490-500G; prepared by manufacturers’ protocol), Difco R2A agar (BD Brand Difco R2A Agar, 218263-500G; prepared by manufacturers’ protocol), Zobell marine agar (Himedia, M384-500G; prepared by manufacturers’ protocol), and seawater agar using glass beads (VWR). 1 liter of seawater agar is made by combining 250mL artificial seawater (32 g reef salt (Instant Ocean) in 1 L MiliQ) with 750 mL 0.22 µm vacuum filtered seawater. To this, 10 g beef extract (Sigma-Aldrich, B4888-500G), 10 g peptone (Research Products International, P20240-500.0) and 20 g Bacto-agar (BD Brand Bacto Agar, 214010-454G) were dissolved and autoclaved in a water bath. Agarose was then cooled on a spin-plate and then transferred into sterile 10 cm dishes and finally cooled to room temperature before storage till use at 4 °C.

Petri dishes were then wrapped in Parafilm and incubated at 20°C for 2 days to 2 weeks. Individual colonies of distinct color, shape, and size were isolated and transferred to fresh plates corresponding to their initial growth condition to produce monocultures. Plates were wrapped in Parafilm and incubated at 20 °C until colonies appeared. Isolates were expanded in liquid minimal starch-yeast (SY) medium (1.25 g starch and 1 g yeast extract in 500 mL ND96). Isolate glycerol stocks were generated by mixing liquid cultures with an equal volume of a 2x freezing stock solution (20% DMSO, 30% seawater, 50% glycerol).

Liquid cultures were verified as monocultures by 16S Sanger sequencing of the bacterial pellet, prepared at 4 °C by spinning at 20,000 RCF. DNA extraction and 16S Sanger sequencing were carried out by Azenta/Genewiz. Briefly, 8F and 1492R primers were used to amplify ribosomal RNA V1-V9. Reads were quality trimmed in Geneious using an error probability limit of 0.05. Consensus sequences for isolates were generated through de novo assembly of forward and reverse reads with >40% quality scores. 295 isolates were verified and identified taxonomically by BLAST search against the NCBI 16S ribosomal RNA database.

Initial trees produced from 16s rRNA multiple sequence alignments of all 295 isolates exhibited many polytomic nodes, indicative of high sequence similarity between isolates. To identify genetically distinct isolates and to reduce data redundancy, all Sanger sequences were clustered using a greedy, global alignment clustering algorithm, cd-hit (identity = 95%)^35^. Consensus sequences were generated for clustered groups using MAFFT (v7.490) alignment (200PAM/ k = 2, with a gap penalty of 3 and an offset value of 0.123)^36^. Evolutionary relationships among remaining distinct strains were inferred by constructing maximum-likelihood phylogenetic trees based on the multiple sequence alignment generated by the SILVA ACT: Alignment, Classification and Tree Service tool (SINA 1.2.12) (SSU). The final tree was constructed using randomized axelerated maximum-likelihood (RAxML) with 1000 bootstrap replicates, GTR model, and Gamma rate^37^.

##### Microbiome sequencing and analysis

The microbiome swab samples were processed and analyzed with the ZymoBIOMICS^®^ Service: Targeted Metagenomic Sequencing (Zymo Research, Irvine, CA). Briefly, DNA was extracted with ZymoBIOMICS^®^-96 MagBead DNA kit (Zymo Research, Irvine, CA) for a final elution volume of 50uL. The DNA samples were prepared for targeted sequencing with the *Quick*-16S^TM^ NGS Library Prep Kit using the custom primer set, *Quick*-16S^TM^ Primer Set V3-V4 (Zymo Research, Irvine, CA). The sequencing library was prepared using a library preparation process in which PCR reactions were performed in real-time PCR machines to control cycles and therefore limit PCR chimera formation. The final PCR products were quantified with qPCR fluorescence readings and pooled together based on equal molarity. The final pooled library was cleaned up with the Select-a-Size DNA Clean & Concentrator^TM^ (Zymo Research, Irvine, CA), then quantified with TapeStation^®^ (Agilent Technologies, Santa Clara, CA) and Qubit^®^ (Thermo Fisher Scientific, Waltham, WA). The ZymoBIOMICS^®^ Microbial Community Standard (Zymo Research, Irvine, CA) was used as a positive control for each DNA extraction and library preparation. Negative controls (including blank extraction control and blank library preparation control) were included to assess the level of bioburden carried by the wet-lab process. The final library was sequenced on Illumina^®^ MiSeq^TM^ with a v3 reagent kit (600 cycles). The sequencing was performed with 10% PhiX spike-in.

Unique amplicon sequences were inferred from raw reads using the Dada2 pipeline^38^. Chimeric sequences were also removed with the Dada2 pipeline. Taxonomy assignment was performed using Uclust from Qiime v.1.9.1. Taxonomy was assigned with the Zymo Research Database, a 16S database that is internally designed and curated, as reference. Composition and abundance visualization and beta-diversity analyses were performed with R Studio (version 2022.07.0, Build 548). Beta diversity matrices were produced using the Bray-Curtis dissimilarity index using a rarefied dataset by the avgdist and metaMDS function in vegan (version 2.6-6.1)^39^. Samples were rarefied to match the sampling depth of the least sampled site. LEfSe was performed using Mothur^40^.

Microbiome ASVs and isolates were matched with each other by creating a database of cultured strain V1-V9 sequences and executing a BLAST search. Any ASV that had >99% pairwise identity with a singular isolate belonging to a cluster (see microbiome sampling and isolate cultivation methods) was designated as a match for that entire cluster.

#### Natural Products Chemistry

##### Bacterial culture, extraction, fractionation, and metabolite purification

Small amounts of bacterial colonies were initially inoculated into 5 mL of SY media (containing 10 g of soluble starch and 4 g of yeast extract in 1 L of sterilized artificial seawater) and cultured on a shaker (180 rpm) at 30 ΔC for 2 days. The initial seed culture was then transferred to 2 L of SY media in 2.5 L Ultra Yield flasks for scale-up culture (120 rpm). After 5 days of cultivation at 30 ΔC, 10 g/L of Amberlite XAD-7 resins were added to the flasks and further shaken for 3 h. XAD-7 resins, absorbed with molecules, were collected using cheesecloth and washed with deionized water to eliminate residual media components. Acetone and methanol (3:1 mixture) was used to elute the absorbed molecules and dried under low pressure yielding crude extracts. The dried materials were re-dissolved in HPLC grade of methanol and injected into preparative HPLC coupled with a C_18_ column (Phenomenex, Luna Omega, 250 × 21.2 mm) for fractionation. Bioactive fractions were then further purified using semi-preparative HPLC with a C_18_ column (Phenomenex, Kinetex, EVO-C_18_, 250 × 10 mm). LC/MS chromatograms and low-resolution electrospray ionization mass spectrometry (LR-ESI-MS) data were obtained on an Agilent Technologies 1200 series high-performance liquid chromatography (HPLC) coupled with an Agilent Technologies 6130 series single quadrupole mass spectrometer and an analytical column (Phenomenex, Kinetex, EVO-C_18_, 100 × 4.6 mm).

##### NMR spectroscopy

All the ^1^H, ^13^C, and 2D NMR spectral data including gCOSY (gradient ^1^H−^1^H correlation spectroscopy), gHSQC (gradient ^1^H−^13^C heteronuclear single quantum coherence), and gHMBC (gradient ^1^H−^13^C heteronuclear multiple bond connectivity) were measured using a Bruker Avance NEO 600 MHz NMR spectrometer, and the chemical shift values are represented on a δ scales. Methanol-*d*_4_ (CD_3_OD) was used as NMR solvent, and the residual protium in the solvent was used to reference chemical shifts (δ_H_ 3.31 and δ_C_ 49.0). All the one-bond ^1^H−^13^C correlations were assigned based on ^1^H, ^13^C, and gHSQC NMR data, and the chemical structures were determined by combinatory analysis of gCOSY and gHMBC NMR spectral data along with 1D NMR data. MestReNova software (v.14.3.1) was used to analysis NMR spectra.

##### Structural determinations

1-acetyl-3-carboxy β-carboline (1A3) (**1**) was isolated as a yellow powder, and its molecular formula was determined to be C_14_H_10_N_2_O_3_ based on the HR-ESI-MS data ([M−H]^−^ *m*/*z* 253.0611, calcd for C_14_H_9_N_2_O_3_ = 253.0619). During its ^1^H, ^13^C, and HSQC NMR analysis, ^1^H and ^13^C signals corresponding presence of one amide proton (δ_H_ 12.07), five of olefinic protons (δ_H_ 9.08, 8.39, 7.82, 7.60, and 7.33), and acetyl methyl protons (δ_H_ 2.84 [3H]) in ^1^H NMR spectrum and two of carbonyl carbons (δ_C_ 201.4 and 167.2), eleven of olefinic carbons (δ_C_ 142.2, 134.7, 134.6 [2C], 131.3, 129.0, 122.0, 120.6, 120.4 [2C], 113.2), and a deshielded aliphatic carbons (δ_C_ 25.8) in ^13^C NMR data were detected. All the one-bond ^1^H−^13^C correlations were identified as **Table S2** based on its ^1^H, ^13^C, and HSQC NMR spectral data. Combined analysis of the COSY and HMBC NMR spectral data revealed its β-carboline partial structure, and two- and three-bond ^1^H−^13^C HMBC correlations of H-4 (δ_H_ 9.08)/C-10 (δ_C_ 167.2), H_3_-12 (δ_H_ 2.84)/C-11 (δ_C_ 201.4), and H_3_-12/C-1(δ_C_ 134.6) identified that the additional acetyl- and carboxylic acid groups were substituted at C-1 and C-3 (δ_C_ 134.6), respectively.

The molecular formula of 1-methyl-3-carboxy β-carboline (**2**) (C_13_H_10_N_2_O_2_) was also confirmed based on its HR-ESI-MS data ([M−H]^−^ *m*/*z* 225.0666, calcd for C_13_H_9_N_2_O_2_ = 225.0670), and all the chemical shift values and one-bond ^1^H−^13^C correlations were assigned by HSQC NMR analysis (**Table S2**). Comparative analysis of 1D NMR data of **1** and **2** indicated that this β-carboline possessed one less carbonyl carbon in the downfield region, which meant absence of acetyl group in its chemical structure. In the same way with **1**, 3-carboxy β-carboline moiety was also determined, and ^1^H−^13^C HMBC correlations from H_3_-11 (δ_H_ 2.83) to C-1 (δ_C_ 141.4) and C-9a (δ_C_ 135.9) revealed that presence of aromatic ring-substituted methyl group at C-1 instead of acetyl group of **1**, completing the full planar structure.

Lumichrome (**3**) was initially dereplicated based on its characteristic UV spectrum (λ_max_ = 220, 260, 350, and 384 nm) and HR-ESI-MS data ([M−H]^−^ *m*/*z* 241.0741, calcd for C_12_H_9_N_4_O_2_ = 241.0731). 1D and 2D NMR spectral data of **3** were acquired and its 1,2-dimethyl benzene moiety was constructed. The other fully substituted carbons, including two carbonyl carbons, was assigned based on their δ_C_ values (δ_C_ 160.6, 150.1, 146.4, and 130.2) and ^1^H−^13^C HMBC correlations from amide protons (δ_C_ 11.84 and 11.67).

##### Natural product environmental detection

“Live crab” samples were pithed and then harvested. For “decay crab” samples, crabs were pithed and then allowed to decay in seawater in a glass beaker covered in parafilm at room temperature for 24 hours. Only female crabs of similar size were used for this analysis. To process samples, crab carapaces and pleons were removed from the body. The internal face of these exoskeleton parts was cleaned thoroughly with a cotton swab to remove all visible biological material. Shells were then placed in a scintillation vial and lightly crushed with 500 µL of freshly prepared 100% MeOH. Vials were vortexed for 1 min and then allowed to sit for 15-30 min. The liquid extract was moved to an Eppendorf and spun at 13,500 rcf for 2 min, to pellet any debris. The supernatant was removed, diluted 1:1 with 80% MeOH.

Eggs were collected at a maximum of 12-24 hours following clutch ejection or removed from an active clutch by hand and then frozen at -80 ΔC till processing. Eggs were desiccated overnight in a lyophilizer and then crushed with a pestle. 200uL of fresh 80% MeOH was added to each sample. All samples were filtered through a 0.45 µM filter (MSRLN04, Millipore Sigma) by spinning at 700 rcf for 2 min before LCMS analysis.

##### LC/MS-QTOF detection and analysis

Samples were analyzed on an Agilent 1290 Infinity II UHPLC coupled to an Agilent 6546 Q-TOF mass spectrometer. An Agilent ZORBAX RRHD Eclipse Plus C18 column (2.1 × 50 mm, 1.8 μm) was used for LC separation with water + 0.1% formic acid (Solvent A) and acetonitrile + 0.1% formic acid (Solvent B) as mobile phases. A linear gradient of 3-50% solvent B was administered over the course of 5 min at a flow rate of 0.6 mL min^-1^. All samples were analyzed using electrospray ionization (ESI) in positive ionization mode. Environmental samples were compared to authentic chemical standards. LC/MS data was analyzed using MassHunter Qualitative Analysis software (version 10.0).

##### UV-imaging

Samples were illuminated with the long-wave UV light setting of a Mineralight Lamp (Model UVGL-58, 366 nm). Standards were prepared in 10% DMSO. Photos of eggs and LUM were acquired with a Nikon D3200 and an AF-S Micro NIKKOR 4mm 1:2.8G lens outfitted with a Tiffen 15 filter. Photos of crabs and H3C were acquired with an iPhone XR.

#### Electrophysiology

##### Xenopus oocyte injection and two-electrode voltage clamp

Defolliculated oocytes were purchased from Ecocyte or Xenopus1 and stored in modified Barth’s solution (88 mM NaCl, 1 mM KCl, 5 mM Tris-HCl, 1 mM MgSO_4_, 0.4 mM CaCl_2_, 0.33 mM Ca(NO_3_)_2,_ 2.4 mM NaHCO_3_, pH 7.4) supplemented with 0.1 mg ml^−1^ gentamycin for up to one week at 4 °C until use. For oocyte expression, plasmids were linearized using Not1-HF (catalog no. R3189, New England Biolabs) for 2 h at 37 °C. Linearized DNA was purified using a PCR purification kit (catalog no. 28104, QIAGEN) and eluted in 30 µl RNase-free water. RNA synthesis was performed with 1–3 µg DNA using the mMESSAGE mMACHINE T7 Transcription Kit including 15 min of DNase treatment (catalog no. AM1344, Ambion). RNA was treated with a Zymo Clean & Concentrator Kit. Oocytes were injected with 2-10 ng RNA using a Nanoject III (Drummond Scientific) and incubated in modified Barth’s solution at 17 °C overnight. Two-electrode voltage clamp recordings were carried out at room temperature with an Oocyte Clamp OC-725C amplifier (Warner Instruments) and digitized using a Digidata 1550B interface and pClamp 11 software. Data were filtered at 1 kHz and sampled at 10 kHz. Recordings were performed using borosilicate glass pipettes with resistances of 1–2 MΩ when filled with 3 M KCl. All chemicals were diluted in ND96 extracellular solution (96 mM NaCl, 2 mM KCl, 5 mM HEPES, 1 mM MgCl_2_, 2 mM CaCl_2_ adjusted to pH 7.4 with NaOH). Stimuli were delivered using a micropipette positioned in the bath. Stimulus-evoked currents were obtained using 200 ms voltage ramps from −120 mV to 80 mV applied every 500 ms withan interstimulus holding potential of −40 mV. For all recordings, responses were assessed as the average current elicited during a 20 ms window, 40 ms after the start the ramp (roughly between -116 and -108 mV). Values were normalized to either the response to 35 µM nootkatone (for CRT1 expressing and uninjected oocytes) or to 10 µM chloroquine (for CR840 expressing oocytes).

To prepare supernatants from bacterial liquid cultures for oocyte screening, cultures were grown from a 1 mm^3^ glycerol stab in minimal starch-yeast media for 8 days at 22 °C. Bacterial cultures were pelleted by centrifugation at 20,000 rcf. Supernatants were filtered for small molecules by centrifugation at max speed through a 3 kDa Amicon Ultra centrifugal filter (catalog no. UFC500324, EMD Millipore). Filtrates were pH-corrected to 7.4 using 2 M HEPES prepared in MilliQ H_2_O. To screen HPLC-separated subfractions, lyophilized samples were dissolved with 9% DMSO/MilliQ H_2_O solution and then diluted to 6.25-25 ng/uL with ND96 for screening.

##### HEK cell culture, transfection and whole-cell voltage clamp

HEK293 cells (authenticated and validated as negative for *Mycoplasma* by the vendor ATCC) were cultured in Dulbecco’s modified Eagle’s medium (DMEM) (Gibco) supplemented with 10% FBS (Peak Serum, US Origin) and 50 IU ml^−1^ penicillin and 50 μg ml^−1^streptomycin (Gibco) at 37 °C and 5% CO_2_ using standard techniques. For transfection, HEK 293 cells were washed with Opti-MEM Reduced Serum Medium (Gibco) and incubated with transfection mix containing 1 µg plasmid CR DNA, 0.3 µg green fluorescent protein plasmid DNA, and 4 µl Lipofectamine 3000 Transfection Reagent (Invitrogen) in Opti-MEM for 4-8 h at 37 °C. Cells were then replated onto glass coverslips in DMEM, incubated for 2 h at 37°C and then moved to 30 °C incubation overnight. Mutagenesis of pUNIV-ObCRT1 to create point mutants and epitope-tagged variants was performed by GenScript.

Patch clamp recordings were carried out at room temperature using a MultiClamp 700B amplifier (Axon Instruments) and digitized using a Digidata 1550B (Axon Instruments) interface and pClamp software (Axon Instruments). Whole-cell recording data were filtered at 1 kHz and sampled at 10 Hz. Voltage-gated currents were leak-subtracted online using a p/4 protocol, and membrane potentials were corrected for liquid junction potentials. For whole-cell recordings in HEK 293 cells, pipettes were 1–3 MΩ. The standard extracellular solution contained): 140 mM NaCl, 5 mM KCl, 10 mM HEPES, 2 mM CaCl_2_, 2 mM MgCl_2_, pH 7.4. The intracellular solution contained: 140 mM Cs^+^ methanesulfonate, 1 mM MgCl_2_, 5 mM NaCl, 10 mM CsEGTA, 10 mM HEPES, pH 7.4.

In ion substitution experiments, relative permeability was determined after the substitution of equimolar monovalent cations by measuring the shift in E_rev_ from I-V relationships filtered at 25 Hz. With each solution exchange, reversal potentials exhibited time-dependent variation so reversal potentials were only assigned when a steady-state was reached. Since NOR and LUM exhibit distinct efficacies for CRT1 activation, ligand concentrations were chosen to be equi-active with respect to the amount of voltage-dependent basal current at a membrane potential of -115mV: 250 ΔM LUM or 100 ΔM NOR. The extracellular solution contained (mM) 150 Na^+^, Cs^+^, or 100 Ca^2^ and intracellular solution contained 150 CsCl and 1 CsEGTA. Solutions were buffered with 10 mM HEPES. Permeability ratios were estimated using the Goldman-Hodgkin-Katz (GHK) equation: P_Na_/P_Cs_ = (γ_Cs_[Cs^+^]_Cytoplasmic_/γ_Na_[Na^+^]_Luminal_)(exp(E_rev,Na_-E_rev,Cs_)F/RT)); P_Ca_/P_Cs_ = γ_Cs_ [Cs^+^]_Cytoplasmic_/4γ_Cs_[Ca^2+^]_Luminal_)(exp(E_rev,Ca_-E_rev,Cs_)F/RT)). γ_Na_=0.75; γ_Cs_=0.71; γ_Ca_=0.29.

Perfusion experiments were performed using a SmartSquirt Micro-Perfusion system (Automate Scientific) pressurized to approximately 30 kPa. All chemical agonists were dissolved in 1% DMSO and were applied until current response reached steady state. Agonists were washed off with ringer until the cell base line returned to its basal state. Supernatants from bacterial liquid cultures were similarly applied to HEK293 cells via perfusion. Agonist-evoked currents were measured as the average current elicited during a 20 ms window, 40 ms after the start of a 500 ms ramp from −120 to 80 mV (roughly between -120 and -112 mV). Current amplitudes were normalized by the amount of basal current present in the absence of ligand. For concentration-response experiments, curves were fit with Hill curves in PRISM to extract EC50 and top values, as many ligands exhibited partial agonism. Correlations with chemical metrics were based on parameters from Chemicalize and PubChem.

#### Structural Biology

##### CRT1 expression and purification

CRT1 was subcloned into the pSBtet-GP (pSBtet-GP-CRT1) vector and appended with a serine linker followed by a single Strep-tag at the C-terminus for affinity chromatography. The pSBtet-GP vector was a gift from Eric Kowarz (Cat # 60495, Addgene)^41^. The Sleeping Beauty transposase system was used to generate a stable cell line for CRT1 expression. HEK293s GnTI^−^ cells (ATCC CRL-3022) were transfected with plasmid DNA (pSBtet-GP-CRT1 + pCMV(CAT)T7-SB100) and cultured in DMEM supplemented with 10% FBS at 37 °C and 8% CO_2_. The pCMV(CAT)T7-SB100 plasmid was a gift from Zsuzsanna Izsvak (Cat # 34879, Addgene)^42^. After 24 hr, 1 μg/mL puromycin was added to select for cells stably expressing CRT1. Selection was carried out for 9 days and terminated after 90% of the cells showed green fluorescence. After confirming the expression of the target gene, cells were adapted to suspension culture in Freestyle293 media (Thermo) supplemented with 2% FBS and penicillin streptomycin (Gibco). They were grown to 6.4 L at a density of approximately 3.5 - 4.0×10^6^ cells per milliliter at 37 °C. Then, 1 μg/mL doxycycline and 1 mM sodium butyrate were added to induce the expression of target protein and cells were cultured at 30 °C. Cells were harvested 48 hr after induction by centrifugation, resuspended in TBS (20 mM Tris, 150 mM NaCl, pH 7.4) containing 1 mM phenylmethanesulfonyl fluoride (Sigma-Aldrich), then lysed using an Avestin Emulsiflex. Lysed cells were centrifuged for 15 min at 10,000 g and the supernatant was collected and centrifuged again for 2 hr at 186,000 g. The membrane pellet was stored at -80 °C until use.

The membrane pellet was thawed in TBS + 1 mM PMSF and mechanically homogenized by a Dounce homogenizer. The membrane was solubilized for 1 hr at 4 °C in TBS buffer with 40 mM *n*-dodecyl β-D-maltoside (DDM, Anatrace). Solubilized membranes were centrifuged for 40 min at 186,000 *g* at 4 °C. The supernatant was collected and passed over Strep-Tactin affinity resin (2 ml) at 0.8 ml/min. The resin was washed with TBS buffer containing 1 mM DDM, then eluted with elution buffer (20 mM Tris, 150 mM NaCl, pH 8.0) supplemented with 5 mM D-desthiobiotin (Sigma-Aldrich) and 1 mM DDM. The CRT1 protein was concentrated to ∼500 μL for nanodisc reconstitution.

##### Nanodisc reconstitution

The plasmid for saposin A expression was obtained from Salipro Biotech AB^43^. Reconstitution was performed by modifying an established protocol^44^. Saposin and soy polar lipids were mixed and nutated at 4 °C for 1 hr, then protein and ligands (0.5-1 mM NOR, H3C and LUM in 1%-2% DMSO) were added and nutated for 30 min at 4 °C. The molar ratio of protein, saposin and lipids was 1:50:250. 100 mg bio-beads were added to remove the detergent at 4 °C overnight. Another 50 mg bio-beads were added the next day and incubated for 2 hr. The receptor-lipid disc mixture was assayed by tryptophan fluorescence SEC and concentrated, then centrifuged for 20 min at 40,000 rpm to remove aggregates before size-exclusion chromatography (SEC). The resulting supernatant was separated by SEC using a Superose 6 Increase 10/300 GL column (Cytiva) equilibrated in TBS buffer at pH 8.0 containing 250-500 μM corresponding ligands in 1%-2% DMSO. SEC fractions corresponding to pentameric CRT1-agonist were pooled and concentrated and the quality of final sample was assessed by tryptophan fluorescence SEC before freezing grids.

##### Cryo-EM sample preparation

For CRT1-NOR, CRT1-H3C and CRT1-LUM, a total of 3 μL of concentrated sample was applied to copper R1.2/1.3 300 mesh holey carbon grids (Quantifoil) that were glow-discharged at 30 mA for 80 s. The grids were immediately blotted for 2.5 s under 100% humidity at 4 °C and then plunge-frozen into liquid ethane cooled by liquid nitrogen using a Vitrobot Mark IV (Thermo Fisher Scientific).

##### Cryo-EM data collection and data processing

For CRT1-NOR complexes, dose-fractioned images were collected on a 300 kV Titan Krios G4 (Thermo Fisher) at UCSD. Images were recording on a Falcon 4 direct electron detector with Selectris X energy filter. The total exposure was 50 e^-^/ Å^2^ and the defocus range was set to -1.4 μm to -2.4 μm with a magnification of 130,000× and 0.935 Å per pixel. For CRT1-H3C and CRT1-LUM, dose-fractioned images were collected on a 300 kV Titan Krios G3 (Thermo Fisher) at UCSD, which has a Gatan K3 direct electron detector and Gatan Biocontinuum Energy Filter. The total exposure was 50 e^-^/Å^2^ and the defocus range was set to -1.4 µm to -2.2 µm with a magnification of 81,000× and 1.072 Å per pixel. The number of micrographs for each data set ranged between 3,300 and 14,600 (Table S4).

All data processing was done using cryoSPARC v.4.4 and v.4.5^45^. Dose-fractionated images were gain-normalized and motion corrected in a CryoSPARC live session with default settings. The contrast transfer function was calculated through patch CTF. Particles were picked by template picker with a diameter of 200 Å and extracted with a 320 pixel box size. After several rounds of two-dimensional (2D) classification, ab-initio and heterogeneous refinement were used to filter the junk particles. Selected particles were subjected to non-uniform (NU) refinement to generate the initial 3D volume with C5 symmetry^46^. To better identify the density of ligands in each pocket, symmetry expansion was performed with C5 symmetry. For each group, particle subtraction was performed to generate the particles with only a single ECD subunit interface, followed by a 3D classification with 10 classes. Particles in each 3D class were re-extracted to full receptor volumes and subjected to NU refinement to generate a 3D map. Classes with ligand density in the pocket were selected and grouped together followed by removing duplicate particles. Finally, particles with density in at least four pockets were grouped and NU refinement with C5 symmetry was used to generate the final maps for CRT1-Norharmane (43,339 particles at a resolution of 3.13 Å), CRT1-H3C (60,658 particles at a resolution of 3.04 Å) and CRT1-Lumichrome (44,653 particles at a resolution of 3.28 Å), respectively.

##### Model building, refinement and validation

PDB: 8EIS was used as the starting model^15^. The starting model was aligned and fitted into the EM density map in UCSF ChimeraX (v.1.7.1)^47^, followed by several rounds of manual building in Coot (v.0.9.8.93) and global real space refinement in Phenix (v.1.20.1) using secondary structure restraints and Ramachandran restraints^48,49^. Model geometry and clash scores were checked by Molprobity^50^. The ligand restraint files were generated using the Grade Web Server (https://grade.globalphasing.org/) with default settings. Figures of protein structures and density maps were generated using UCSF ChimeraX and PyMOL (v.2.5.5, Schrodinger, LLC). The pore radius profiles were calculated by HOLE v.2, and the hydrophobicity of the permeation pathway was generated by CHAP^51,52^. The 2D ligand interaction diagram was analyzed by Schrodinger Maestro. Programs were compiled by SBGrid^53^.

#### Molecular evolution

To evaluate the extent of selective pressure acting on ligand-binding sites, likelihood ratio test values from Allard and Kang et al., 2023 were used^15^. Briefly, codon alignments of CR sequences were used to estimate the ratio of synonymous and non-synonymous substitutions (*ω*) across AChR-like genes. The likelihood of a single *ω* across branches (M0 model), branch-specific *ω* (free ratio), and distinct *ω* partitioning in the CR clades was used as foreground and the remaining AChR-like genes was a background (two-ratio). We also tested partitioning *ω* between the two major CR clades (three-ratio). Initial *ω* values were varied to check for convergence and likelihood ratio tests were used to determine the best-fitting models and sites under stronger diversifying selection. To further characterize positive selection across sites in the CR clades, we used the more sensitive and robust MEME model from HyPhy, with a default threshold of significance of LRT = 3 for positively selected sites^54^.

#### Ex vivo analyses

##### Axial nerve recording

Arm tips from sedated animals were transferred to a 10 cm Petri dish containing 20 mL holding solution (430 mM NaCl, 10 mM KCl, 10 mM HEPES, 10 mM CaCl_2_, 50 mM MgCl_2_, 10 mM D-glucose, pH 7.6). Nerve recordings were performed using a borosilicate glass suction electrode shaped and polished to fit over the entire cut end of the radial nerve. A high resistance reference electrode was placed in the bath. 1 mL extracts of 3 kDa-filtered natural product (50 µL) or ligand (1 mL, prepared in 1-10% DMSO), were applied to the tip of the arm at about 1.5 cm distance. Experimenters were blinded to the identity of the chemical during data acquisition and analysis. The order of the stimuli applied to the arm was randomized.

Gap-free recordings were made with 10-kHz sampling at 10,000× gain and signals were high pass-filtered at 100 Hz and lowpass-filtered at 1 kHz using a Warner DP-311A headstage and AC/DC amplifier (Warner) and digitized using a Digidata 1440A digitizer (Molecular Devices) with ClampEx software (Molecular Devices). Recordings were processed using Igor (Igor WaveMetrics v8.04). For quantification of responses, the baseline signal was subtracted, and the root-mean-square of the integrated response was assessed over a 10s window following stimulus or vehicle addition. Squared versions of raw nerve signals are presented for visualization in figures.

Extracts from rejected eggs, mothered eggs, decay crab shells, and freshly-killed crab shells were prepared by redissolving, pulverized samples previously frozen with liquid N_2_. Redissolved samples were filtered by centrifugation at max speed through a 3 kDa Amicon Ultra centrifugal filter (catalog no. UFC500324, EMD Millipore).

##### Semi-autonomous arm movement

Arm tips from sedated animals were transferred to a 10 cm Petri dish containing 50 mL holding solution (and held in place using suction through an appropriately sized black tygon tubing). Ligands were first dissolved in DMSO and then diluted in holding solution to 1%-10% final DMSO; 1 mL stimulus at the indicated concentration was perfused over the arm over 2 s using a micropipette. Arm behavior was recorded at 500 ms intervals using a FLIR grasshopper camera (Teledyne) equipped with a 35 mm Nikon DX AF-S NIKKOR 1:1.8G lens controlled using the Spinnaker SDK software (Teledyne). Videos were segmented and motion was measured using ImageJ (NIH) by measuring Z-stack projections after performing stack difference analysis. Total arm motion in response to each chemical was measured by summing the stack difference and values were normalized such that the response to 1 mM nootkatone was set to 1 for each arm. Experimenters were blinded to the identity of the chemical during data acquisition and analysis. The order of the stimuli applied to the arm was randomized.

##### Octopus sucker cup adhesion

Arm tips from sedated animals were mounted on a glass slide (VWR) using cyanoacrylate glue (SeaChem) with the suckers facing upwards. The glass slide was placed in a 10 cm petri dish containing 50 mL seawater filtered with a 0.22 μm vacuum filter. Separately, glass coverslips were prepared by adding 500 μL of 1% agarose (VWR, N605-500G) infused with chemical to the top of the coverslip. These coverslips were inverted and pressed against the mounted arm tip using forceps, and then removed after 2 s of contact. The assay was recorded at 10 Hz using a FLIR grasshopper camera (Teledyne) equipped with a 35 mm Nikon DX AF-S NIKKOR 1:1.8G lens controlled using the Spinnaker SDK software (Teledyne). Videos were analyzed by manually counting the proportion of total sucker cups that remained attached to the cover slip when lifted 3 mm vertically away from the mounted arm. The coverslips used in this assay were blinded and randomized to the experimenter both while performing the assay and during analysis. The order of the stimuli applied to the arm was randomized.

#### Animal Behavior

##### Octopus microbiome-covered floors

Swabs from living crab carapaces or decayed crab carapaces were used to inoculate glycerol stocks that were preserved at -80 °C until use. Seawater agar plates were inoculated with 50-200 μL of the stock and then grown till lawn formation. Decay crab communities grew faster on seawater agar plate than living crab communities, so fully confluent plates were parafilm sealed and stored at 4 °C until use. Once both communities covered comparable areas, the solid media was removed from the petri dish and laid onto the bottom of an acrylic behavior tank. Cultures were sealed to the bottom by the addition of hot, 1.5% agarose prepared with seawater. Animals were introduced to the tank and allowed to freely explore for the 10 minute trial. The trials were monitored with a GoPro HERO7 camera (GoPro Inc.). Afterward, the animal was returned to its home tanks for at least 24 hours. The number and duration of each arm touch on a plate was characterized during playback in BORIS^55^. Experimenters were blinded to the identity of the community type during data analysis. The placement of the agar disc was randomized for each trial.

##### Octopus crab predation

Octopus were moved to behavior tanks for filming behavior with chemical-doped crab mimics (B00362OM12, Amazon). Animals were acclimated in a behavior tank for the10 min prior to trial start. Each trial lasted 60 minutes. A cohort of 6 animals were used to collect the data, with animals receiving a certain stimulus more than once. Trails were conducted on an animal that had not been fed or moved to the behavior tank in 24 hours immediately preceding. To incentivize interaction with the introduced item, octopus were offered an alternating regimen of mimics and freshly immobilized fiddler crabs. Prior to a trial, mimics were added to beakers containing, 10 mM nootkatone, 10% DMSO, 1 mM H3C, or 1 mM NOR and agitated on a rocker for 25 min. Octopus behavior was analyzed post hoc in playback. The latency of first investigation was the time between crab introduction and the first touch. Engulfing behavior was documented if the octopus ensnared the crab within its brachial web. Trials from octopus that did not interact with the crabs were discarded. Experimenters were blinded to the identity of the chemical of the crab mimic during data acquisition and analysis. For comparisons of octopus predation on freshly killed and 24 hr decayed crabs, octopus were similarly acclimated and filmed.

##### Octopus brooding behavior

False eggs were created in two stages. First, a 1% agarose (VWR, N605-500G) solution was made containing either a chemical stimulus or a vehicle control. 200 μL of agarose was pipetted onto a sheet of Parafilm and left to cool into a half-sphere. Once cooled, the half-sphere was turned flat-side-up and an additional 200 μL of agarose was added to the flat side to create the spherical final false egg shape. Female octopuses that were brooding eggs were recorded with a GoPro HERO7 (GoPro Inc.) while the false agarose eggs were dropped into the animal’s den from above. Trials were recorded until the false egg was rejected from the den, and the time of rejection was noted. Experimenters were blinded to the identity of the chemical during data acquisition and analysis. The order of the stimuli offered was randomized.

