## Supplemental Figures and Tables for "Environmental microbiomes drive chemotactile sensation in octopus"

### Supplemental Figure 1

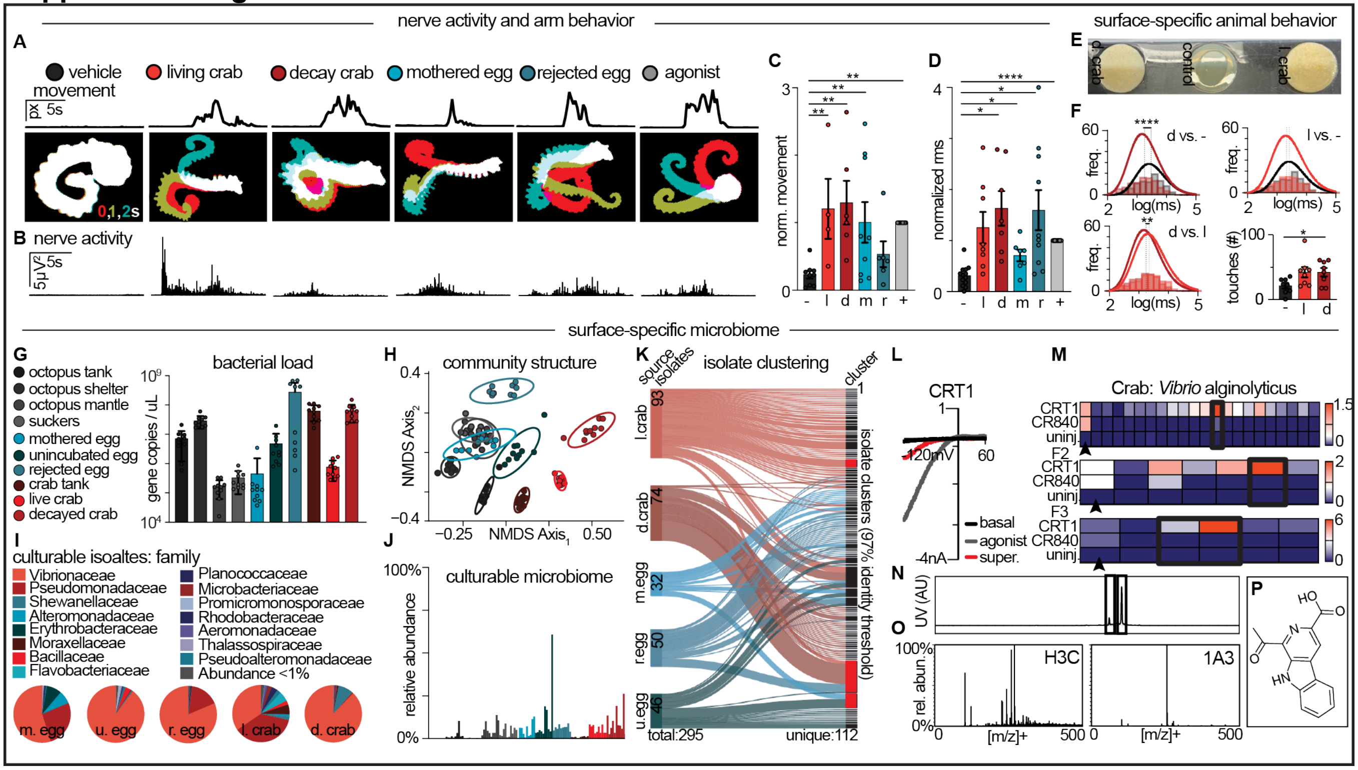

**Supplemental Figure 1 | Microbes produce chemical cues that activate octopus "taste by touch" chemotactile receptors. A**, Octopus arms exhibit autonomous movement in response to <3 kDa-filtered extracts from natural products or a positive control agonist (nootkatone, grey), but not vehicle control. *Top*: change in thresholded pixel values over time. *Bottom*: corresponding arm kymographs, red: 0 s; green: 1 s; teal: 2 s. **B**, Octopus arm axial nerve innervating suckers also responded to these extracts but not to vehicle control. **C,** Quantified arm movement in response to extracts of live (immobilized) crab (l., p = 0.0072), decayed crab (d., p = 0.0011), incubated egg (i., p = 0.0084) and control agonist (nootkatone: p < 0.0089) but not vehicle control incited arm movement by Mixed-effects analysis with Dunnett's multiple comparisons test, with a single pooled variance, n = 4-9 arms. **d,** Quantified nerve responses to extracts of d.crab (p = 0.0406), m.egg (p = 0.0269), r.egg (p = 0.0348) and control agonist (nootkatone: p < 0.0001) but not vehicle control incited axial arm activity by Mixed-effects analysis with a Geisser-Greenhouse correction and Dunnett's multiple comparisons test, with individual variances computed for each comparison, n = 7-13 arms. **E,** Swabs from (L-R) decaying crab and living crabs struck onto agar grown into communities with distinct morphologies. **F,** Octopuses freely explored a tank with inlaid bacterial communities (*E*) and exhibited an increased number of short duration touches on surfaces with decaying crab microbiomes. Touch duration is represented as the geometric mean (dotted lines) extracted from lognormal fits (solid lines): d.crab = 3.246 ms, l.crab = 3.380 ms, negative control = 3.474 ms. Differences in duration were statistically assessed by the arithmetic mean of all touches from 8 trials and were evident in control (3.408 ms) versus d.crab (3.216 ms) (p < 0.0001) and l.crab (3.334 ms) versus d.crab (p = 0.0021) comparisons, but not l.crab versus control by ordinary one-way ANOVA with Tukey's multiple comparisons test, with a single pool variance. The number of touches for the d.crab community was greater than control (p = 0.0315) by a repeated measures, one-way ANOVA with the Geisser-Greenhouse correction for unequal variance with Tukey's multiple comparisons test with individual variances for each comparison. **G,** Quantitative V3-V4 sequencing showed that different surfaces carry different relative abundances of bacteria (p < 0.001, ordinary one-way ANOVA)[34]. **H,** Nonmetric dimensional scaling analysis and Permutational multivariate analysis of variance (PERMANOVA) tests show that these communities have distinct structures that are correlated with the location of sample site. (Bray-Curtis index, F= 20.57624, p < 0.001, see *Table S1* for statistics). **I**, Bacterial strains of diverse families were isolated from five different surfaces in the octopus's environment. **J,** By matching the V1-V9 sequences with the V3-V4 microbiome data, we estimated that <10% of the total microbiome for each surface was cultured.

Key in Fig. 1SG. **K**, To reduce redundancy isolates were clustered into groups using a ~97% identify threshold and from 295 total isolates, 112 clusters were identified as unique. Clusters highlighted in red represent the isolates used for this study. From top to bottom: *Pseudomonas alcaligene*s, *Vibrio alginolyticus*, and *Vibrio mediterranei.* **L**, A *Vibrio alginolyticus* isolate that showed activity in HEK293 cells and in oocytes screens (*Fig. 1D*) was also sub-fractioned and screened for molecules that activated CRT1 using TEVC. **M,** Fractions with the greatest activity were further fractionated and screened. **N**, UV chromatographs of the F3 round of subfractionation revealed two active components. **O**, Mass spectrometry analysis reveals major mass ions of 227.2, (same as *Fig. 1I*) and 255.1. **P,** 1-acetyl-3-carboxy acid β-carboline (1A3) was the major molecular component of the active subfraction from *Vibrio alginolyticus*.

### Supplemental Figure 2

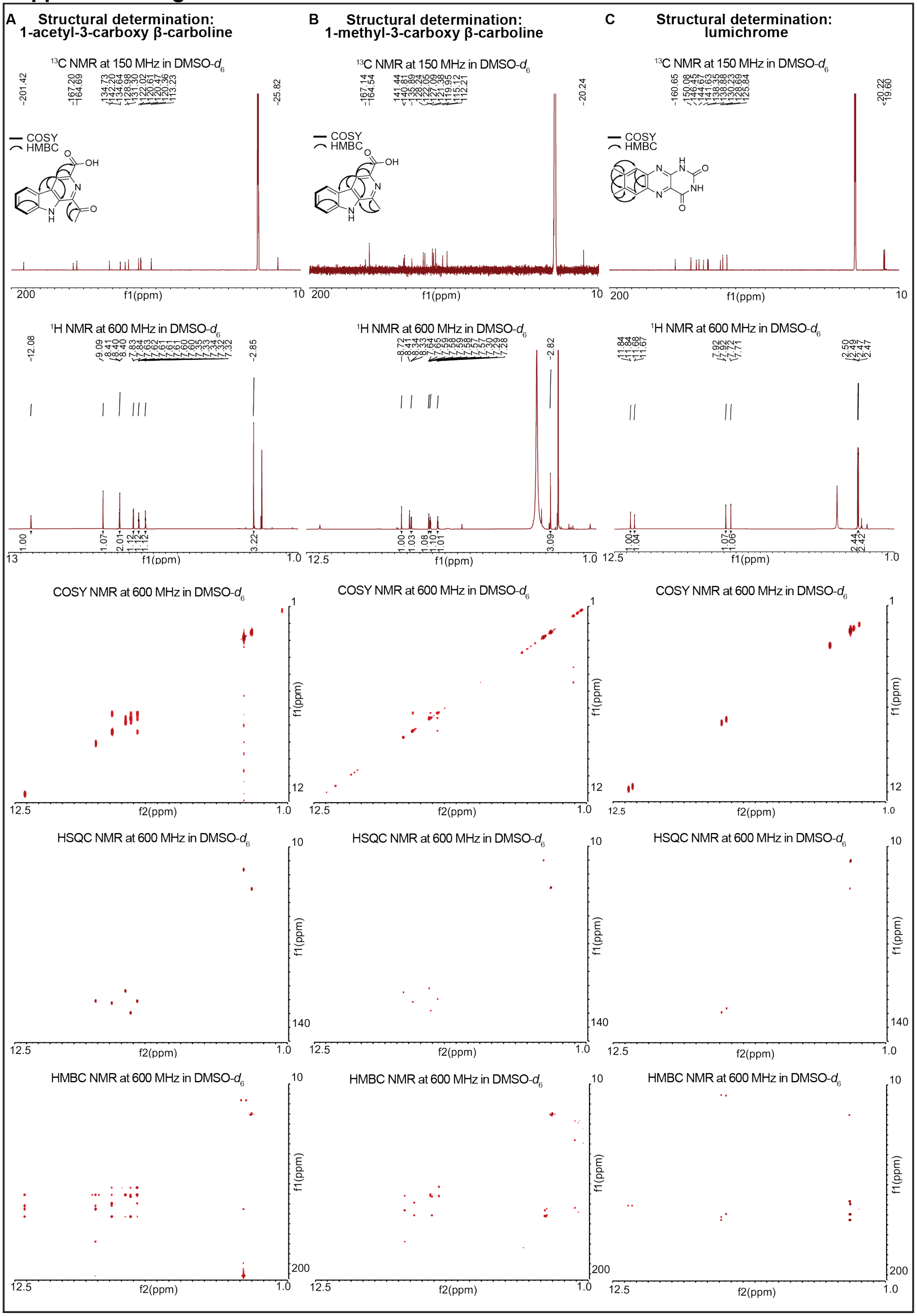

**Supplemental Figure 2 | Structural determination of chemical constituents in active subfractions of bacterial supernatants. Column A,** 1-acetyl-3-carboxy β-carboline (*1*) was isolated as a yellow powder, and its molecular formula was determined to be $C_{14}H_{10}N_2O_3$ based on the HR-ESI-MS data ($[M-H]^-$ *m/z* 253.0611, calcd for $C_{14}H_9N_2O_3$ = 253.0619). During its $^1H$, $^{13}C$, and HSQC NMR analysis, $^1H$ and $^{13}C$ signals corresponding presence of one amide proton ($\delta_H$ 12.07), five of olefinic protons ($\delta_H$ 9.08, 8.39, 7.82, 7.60, and 7.33), and acetyl methyl protons ($\delta_H$ 2.84 [3H]) in $^1H$ NMR spectrum and two of carbonyl carbons ($\delta_C$ 201.4 and 167.2), eleven of olefinic carbons ($\delta_C$ 142.2, 134.7, 134.6 [2C], 131.3, 129.0, 122.0, 120.6, 120.4 [2C], 113.2), and a de-shielded aliphatic carbon ($\delta_C$ 25.8) in $^{13}C$ NMR data were detected. All the one-bond $^1H-^{13}C$ correlations were identified as *Table S2* based on its $^1H$, $^{13}C$, and HSQC NMR spectral data. Combined analysis of the COSY and HMBC NMR spectral data revealed its β-carboline partial structure, and two- and three-bond $^1H-^{13}C$ HMBC correlations of H-4 ($\delta_H$ 9.08)/C-10 ($\delta_C$ 167.2), $H_3$-12 ($\delta_H$ 2.84)/C-11 ($\delta_C$ 201.4), and $H_3$-12/C-1($\delta_C$ 134.6) identified that the additional acetyl- and carboxylic acid groups were substituted at C-1 and C-3 ($\delta_C$ 134.6), respectively. **Column B,** The molecular formula of 1-methyl-3-carboxy β-carboline (*2*) ($C_{13}H_{10}N_2O_2$) was also confirmed based on its HR-ESI-MS data ($[M-H]^-$ *m/z* 225.0666, calcd for $C_{13}H_9N_2O_2$ = 225.0670), and all the chemical shift values and one-bond $^1H-^{13}C$ correlations were assigned by HSQC NMR analysis (*Table S2*). Comparative analysis of 1D NMR data of (*1*) and (*2*) indicated that this β-carboline possessed one less carbonyl carbon in the downfield region, which meant absence of acetyl group in its chemical structure. In the same way with (*1*), 3-carboxy β-carboline moiety was also determined, and $^1H-^{13}C$ HMBC correlations from $H_3$-11 ($\delta_H$ 2.83) to C-1 ($\delta_C$ 141.4) and C-9a ($\delta_C$ 135.9) revealed that presence of aromatic ring-substituted methyl group at C-1 instead of acetyl group of (*1*), completing the full planar structure. **Column C,** Lumichrome (*3*) was initially dereplicated based on its characteristic UV spectrum ($\lambda_{max}$ = 220, 260, 350, and 384 nm) and HR-ESI-MS data ($[M-H]^-$ *m/z* 241.0741, calcd for $C_{12}H_9N_4O_2$ = 241.0731). 1D and 2D NMR spectral data of (*3*) were acquired and its 1,2-dimethyl benzene moiety was constructed. The other fully substituted carbons, including two carbonyl carbons, was assigned based on their $\delta_C$ values ($\delta_C$ 160.6, 150.1, 146.4, and 130.2) and $^1H-^{13}C$ HMBC correlations from amide protons ($\delta_C$ 11.84 and 11.67). Consistent with bacterial biosynthesis, H3C and 1A3 (from crabs) can be produced from a tryptophan or a tryptamine precursor by microbial enzymes using Pictet-Spengler chemistry[18,19]. Similarly, LUM (from eggs) is an oxidation product of riboflavin, which is produced by flavin-dependent monooxygenases[20].

### Supplemental Figure 3

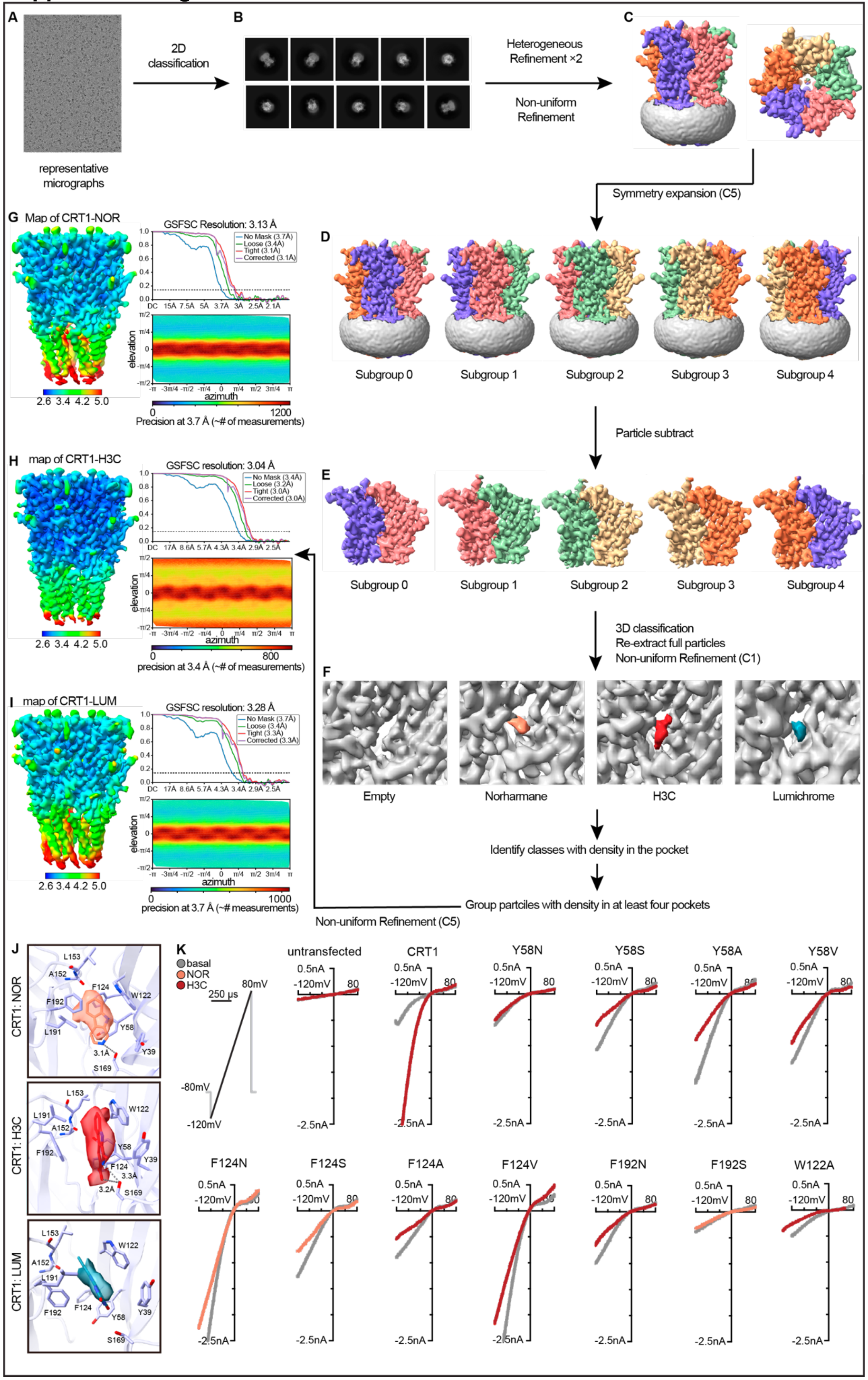

**Supplemental Figure 3 | Cryo-EM Data processing and mutagenesis of octopus chemotactile receptors. A,** Representative cryo-electron micrographs from CRT1-H3C dataset. **B,** Representative 2D classes from the final selected 2D classes of CRT1-H3C complex. **C-F,** Workflow of symmetry expansion and 3D classification to identify particles with density of ligands in each pocket. **F,** Representative classes for an empty pocket or with ligand density in the pocket. **G-I,** Unsharpened map of CRT1-NOR (threshold level of 0.11), CRT1-H3C (threshold level of 0.14) and CRT1-LUM (threshold level of 0.15) colored by local resolution. FSC plot indicates the resolution at FSC=0.143 and angular distribution plot indicates particle orientation preferences. **J,** Orthosteric binding sites for NOR, H3C, and LUM highlight key residues for interaction between ligand and CRT1. **K,** Exemplar basal (grey) and ligand-induced (burgundy or teal) currents from HEK293 cells expressing CRT1 with the indicated ligand-binding pocket mutants. Ligand response was tested using an EC50 concentration of NOR (75 μM) or H3C (300 μM).

**Supplemental Figure 4**

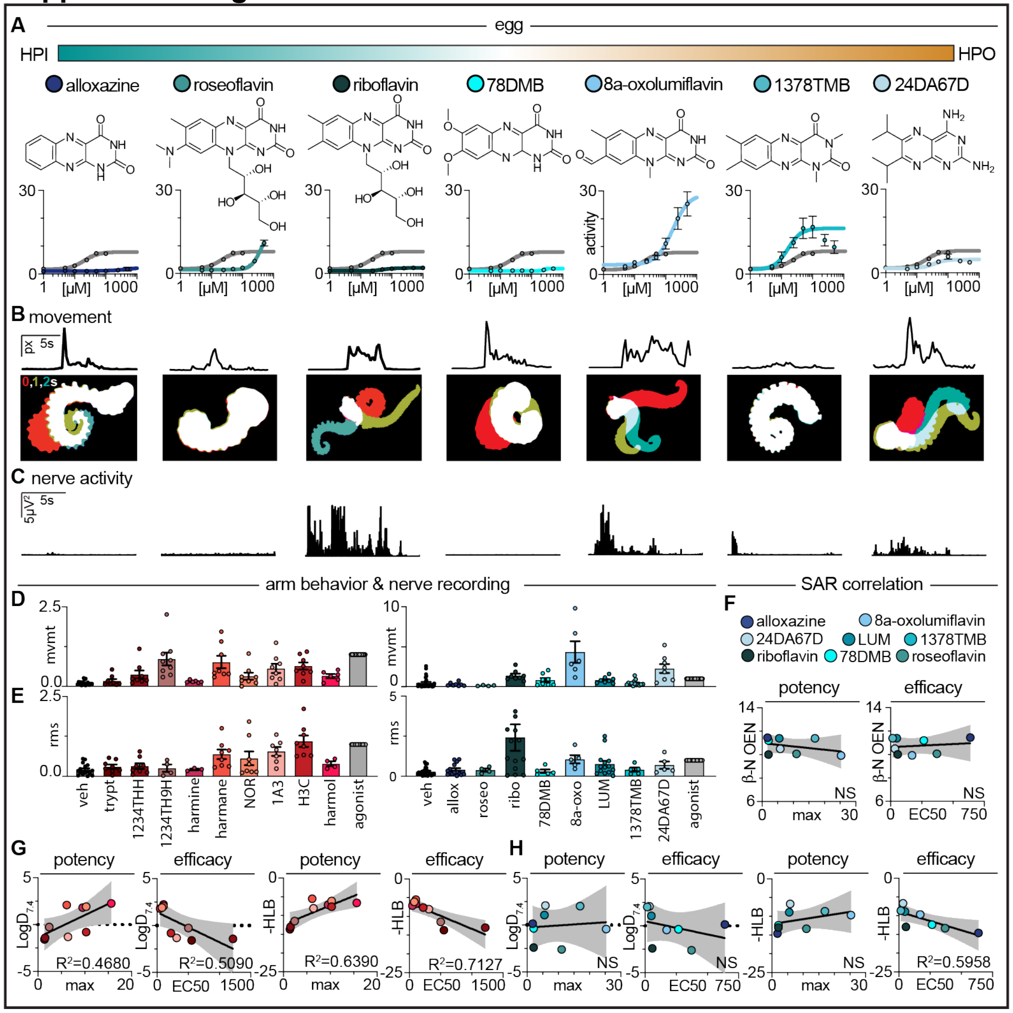

**Supplemental Figure 4 | Analogs of microbial metabolites activate the octopus chemotactile system. A,** Ligand interaction analyses suggest that egg-derived microbial ligands rely heavily on hydrophobic interactions**.** Concentration-response relationships for CRT1 using egg-derived microbial molecules of varying hydrophilic-lipophilic balance (HLC) values. n = 3-8 cells per ligand. See *Table S3* for fit values. **B,** Octopus arms exhibited autonomous movement in response to egg-derived microbial ligands; quantification in (*D*)**.** *Top*: change in thresholded pixel values over time. *Bottom*: corresponding arm kymographs, red: 0s; green: 1s; teal: 2s. n = 4-14 arms. **C,** Axial nerve responses to egg-derived microbial ligands. n = 4-14 arms; quantification in (*D*). **D,** *Left*: 1 mM crab-derived microbial ligands induced arm movement in 1234TH9H (p = 0.0409), H3C (p = 0.0061), and positive control agonist nootkatone (grey) (p < 0.0001). *Right*: 0.5 mM egg-like molecules induced movement in riboflavin (p = 0.0032) and nootkatone (p < 0.0080). 100 µM LUM also induced movement (mean = 0.8470, n = 8, p = 0.004). **E,** *Left*: 1 mM of crab-derived microbial ligands induced nerve activity in harmane (p = 0.0205), H3C (p = 0.0029), 1A3 (p = 0.0404) and nootkatone (p < 0.0001). *Right*: 0.5 mM of egg-derived microbial molecules induced nerve activity in 8a-oxo (p = 0.0391) and nootkatone (p < 0.0001). 100 µM LUM also induced activity (mean = 1.274, n = 14, p = 0.0421). Statistical analyses were computed with a Mixed-effects analysis with Dunnett's multiple comparisons test, with individual variances computed for each comparison. **F,** Linear regression analysis shows that CRT1 activity elicited by egg-derived microbial molecules does not correlate with the orbital electronegativity (OEN) of the 3' nitrogen. See *Table S5* for chemical metrics. **G**, Additional linear regression analyses suggested that crab-molecule activity correlated with LogD (potency: p = 0.0421, efficacy: p = 0.0309) and HLB (potency: p = 0.0097, efficacy: p = 0.0042). **H,** Similar correlations with egg molecules were not as consistent (efficacy: p = 0.0248).

**Supplemental Figure 5**

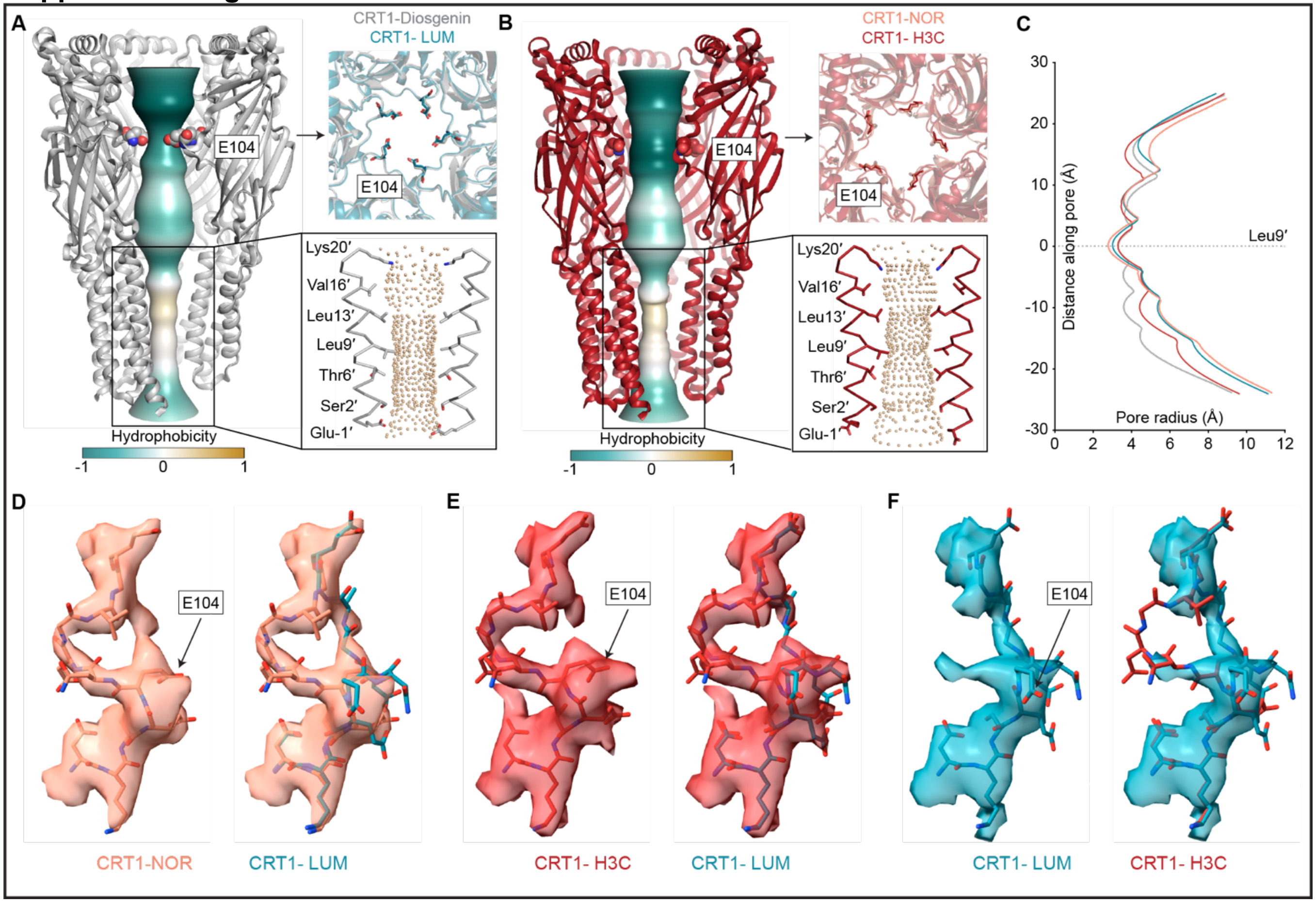

**Supplemental Figure 5 | Different conformations of the Ω-loop and E104 correlate with ligand-dependent differences in relative $Ca^{2+}$ permeability. A,** The ion permeation pathway of the CRT1-disogenin complex (PDB: 8EIS) colored by hydrophobicity. The top view shows a superposition of the LUM and diosgenin-bound structures, highlighting the same conformations of the E104 side chain, correlating with high relative $Ca^{2+}$ permeabilities (*Fig. 6F*). *Inset:* Two M2 helices with pore-lining residues are shown as sticks; spheres indicate pore shape for CRT1-Diosgenin. **B**, Ion permeation pathway of the CRT1-H3C complex colored by hydrophobicity. The top view comparison with CRT1-NOR indicates that E104 adopts a consistent conformation for crab-related molecules. *Inset:* same as (*A*) but for CRT1-H3C complex. **C**, Pore diameters of CRT1-NOR, CRT1-H3C, CRT1-LUM and CRT1-diosgenin indicate that the CRT1 pore is open when crab- and egg-related molecules are bound to the pocket. Leu9' is defined as y = 0. **D**,**E**, Cryo-EM density segment of Ω-loop in CRT1-NOR (threshold = 0.08) and CRT1-H3C (threshold = 0.12), and model of CRT1-LUM was docked into the density, respectively. **F,** Cryo-EM density segment of the Ω-loop in CRT1-LUM (threshold = 0.13), and model of CRT1-H3C was docked into the density. This loop lining the vestibule is remarkably dynamic, adopting ligand-dependent conformations.

### Supplemental Figure 6

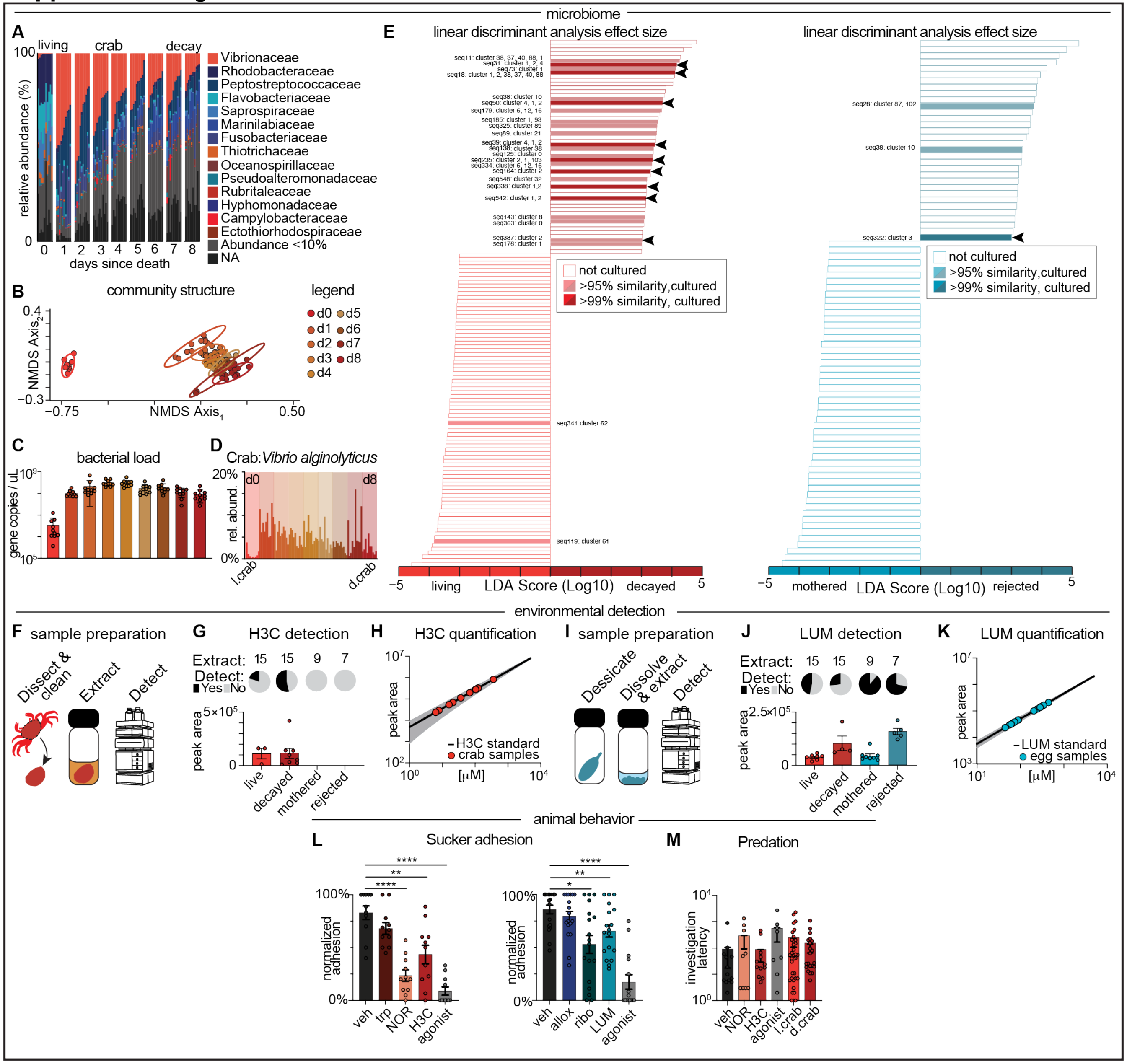

**Supplemental Figure 6 | Microbial metabolites are present in the octopus environment and elicit a behavioral response. A,** Microbiome profiling demonstrates that the communities present on crab carapaces evolve over the time course of decay, with the most striking changes happening less than 24 hrs following death, n = 10. **B,** Nonmetric dimensional scaling analysis and permutational multivariate analysis of variance (PERMANOVA) tests show that the microbial communities on living and decayed crab carapaces have distinct structures. (Bray-Curtis index, F = 13.84954, p < 0.001, see *Table S6* for statistics). **C,** quantitative V3-V4 sequencing suggests that crab carapaces accumulate more copies of the bacterial V3-V4 gene the longer they are left to decay until (d0 vs. d2-d4: p = <0.0001; d0 vs. d5: p = 0.0012; d0 vs. d6: p = 0.0003; d0 vs. d7: p = 0.0446; ordinary one-way ANOVA with Dunnett's multiple comparisons test using a single pooled variance, n = 10). **D,** Amplicon sequencing variants (ASVs) that matched in sequence with the H3C-producing *Vibrio alginolyticus* were enriched in the decaying crab microbial community. **E,** LEfSE (Linear discriminant analysis effect sizes) analysis determines which ASVs (seqs) are most likely to explain differences between *Left*: D0 crabs (red) and D1 decaying crabs (burgundy) and *Right*: mothered eggs (aqua) vs. rejected eggs (teal). ASVs were matched with cultured isolates based on >99% (dark) or >95% (muted) sequence identity to determine which defining strains were cultured. Such pairings suggested that the "*Vibrio alginolyticus* isolate"-like ASVs (black arrows) are highly enriched in decaying crab microbiomes while the "*Vibrio mediterrani* isolate"-like ASV is highly enriched in rejected egg microbiomes (black arrows). **F,** Metabolites were extracted from crab shells by soaking whole shells in MeOH and running LC/MS analysis on the extract. **G,** H3C was detected from crab extracts in 3/9 total extractions from non-decayed crabs and 8/15 total extractions from decayed crabs. LUM was not detected in any of the crab extracts. **H,** H3C concentration in a 0.5 mL extraction volume ranged from 6 μM to 360 μM by interpolation against a standard curve (y = 0.8756x + 3.682, $R^2$ = 0.9755). 1A3 was detected in two crab extractions and had an interpolated concentration of 50-200 μM. **I**, Metabolites were extracted from eggs by lyophilizing whole eggs in MeOH and running LC/MS analysis on the extract. **J,** LUM was detected from egg extracts in 8/9 total extractions from mothered eggs and 5/7 total extractions from rejected eggs. LUM was also detected in 7/15 live crab extracts and 4/15 decay crab extracts. **K,** LUM concentration in a 100 μL extraction ranged from 200 μM to 350 μM, by interpolation against a standard curve (y = 0.9508x + 2.802, $R^2$ = 0.9997). **L,** Autonomous arm behavior showed that sucker adhesion to coverslips was decreased with 1 mM NOR (p < 0.0001), 1 mM H3C (p = 0.0084), or 1 mM nootkatone (p < 0.0001) compared to DMSO as a control by repeated measures one-way ANOVA with the Geisser-Greenhouse correction and Dunnett's multiple comparisons test with individual variances computed for each comparison (n = 11 arms). Similarly, when coverslips were doped with metabolite analogs, 500 μM LUM (p = 0.0096), 500 μM riboflavin (p = 0.0226), or 1 mM nootkatone (p < 0.0001), suckers were less likely to autonomously adhere to the agarose compared to the vehicle control by mixed-effects analysis with the Geisser-Greenhouse correction and Dunnett's multiple comparisons test with individual variances computed for each comparison. (n = 15-18 arms). **M,** Hungry octopuses readily investigated crab mimics, immobilized fresh crabs, and decayed crabs. Regardless if the item presented was a chemical-soaked crab (1 mM H3C: n = 12; 1 mM NOR: n = 9; 10 mM nootkatone: n = 8; DMSO: n = 13) or a real crab (immobilized live: n = 35; decayed: n = 21), the octopus took the same amount of time to first touch (Ordinary one-way ANOVA with Tukey's multiple comparisons test using a single pooled variance).

**Supplemental Tables**

**Supplemental Table 1 | PERMANOVA analysis on microbiome Bray Curtis dissimilarity composition.** Data were computed in the R environment using the command "adonis2". Df: degrees of freedom. Sum Sq: Sum of squares. Pr(>F): P-value associated with the F-statistic. P.adjusted: Bonferroni test adjusted P-value.

| PERMANOVA 9999 permutations. Bray Curtis by sample site. | | | | | | |
|---|---|---|---|---|---|---|
| | Df | Sum Sq | R2 | F-value | Pr(>F) | |
| Groups | 9 | 28.16572 | 0.6728524 | 20.56724 | 0.001 | |
| Residuals | 90 | 13.69446 | 0.3271476 | | | |
| Totals | 99 | 41.86018 | 1.0000000 | | | |
| Pairwise comparisons, multiple comparisons corrected with the Bonferroni test | | | | | | |
| | Df | Sum Sq | R2 | F-value | Pr(>F) | P.adjusted |
| o.tank vs. o.shelter | 1 | 3.471630 | 0.5702925 | 23.88896 | 0.001 | 0.045 |
| o.tank vs. c.tank | 1 | 3.555604 | 0.6356391 | 31.40156 | 0.001 | 0.045 |
| o.tank vs. mantle | 1 | 3.082427 | 0.5177545 | 19.32539 | 0.001 | 0.045 |
| o.tank vs. suckers | 1 | 3.492365 | 0.595095 | 26.45488 | 0.001 | 0.045 |
| o.tank vs. u.egg | 1 | 2.106474 | 0.4076391 | 12.38688 | 0.001 | 0.045 |
| o.tank vs. m.egg | 1 | 2.801051 | 0.4540107 | 14.96768 | 0.001 | 0.045 |
| o.tank vs. r.egg | 1 | 3.607561 | 0.5945023 | 26.38989 | 0.001 | 0.045 |
| o.tank vs. d.crab | 1 | 3.811878 | 0.6203977 | 29.41805 | 0.001 | 0.045 |
| o.tank vs. l.crab | 1 | 4.045331 | 0.6981909 | 41.64035 | 0.002 | 0.090 |
| o.shelter vs. suckers | 1 | 2.361570 | 0.4667951 | 15.75813 | 0.001 | 0.045 |
| o.shelter vs. c.tank 1 | 1 | 3.490878 | 0.5966959 | 26.63134 | 0.001 | 0.045 |
| o.shelter vs. l.crab | 1 | 3.912266 | 0.6539758 | 34.01948 | 0.001 | 0.045 |
| o.shelter vs. d.crab | 1 | 3.666124 | 0.5800993 | 24.86728 | 0.001 | 0.045 |
| o.shelter vs. m.egg | 1 | 1.457446 | 0.2831481 | 7.10979 | 0.001 | 0.045 |
| o.shelter vs. mantle | 1 | 2.474106 | 0.436623 | 13.95019 | 0.001 | 0.045 |
| o.shelter vs. u.egg | 1 | 3.172325 | 0.483979 | 16.8823 | 0.001 | 0.045 |
| o.shelter vs. r.egg | 1 | 3.531674 | 0.5593718 | 22.85077 | 0.001 | 0.045 |
| c.tank vs. suckers | 1 | 3.786082 | 0.6410629 | 32.14807 | 0.001 | 0.045 |
| c.tank vs. l.crab | 1 | 3.372187 | 0.6932212 | 40.67419 | 0.001 | 0.045 |
| c.tank vs. d.crab | 1 | 3.826765 | 0.6482987 | 33.1798 | 0.001 | 0.045 |
| c.tank vs. m.egg | 1 | 3.171609 | 0.5047301 | 18.34382 | 0.001 | 0.045 |
| c.tank vs. mantle | 1 | 3.460295 | 0.5695992 | 23.82148 | 0.001 | 0.045 |
| c.tank vs. u.egg | 1 | 3.275634 | 0.5387289 | 21.0226 | 0.001 | 0.045 |
| c.tank vs. r.egg | 1 | 3.828930 | 0.6346416 | 31.26669 | 0.001 | 0.045 |
| mantle vs. suckers | 1 | 0.9633728 | 0.246002 | 5.872742 | 0.001 | 0.045 |
| mantle vs. l.crab | 1 | 3.741857 | 0.6167485 | 28.96654 | 0.001 | 0.045 |
| mantle vs. d.crab | 1 | 3.496075 | 0.5458376 | 21.6334 | 0.001 | 0.045 |
| mantle vs. m.egg | 1 | 0.3812385 | 0.08812146 | 1.739471 | 0.04 | 1 |
| mantle vs. u.egg | 1 | 2.599232 | 0.4167585 | 12.862 | 0.001 | 0.045 |
| mantle vs. r.egg | 1 | 2.986913 | 0.4958287 | 17.70215 | 0.001 | 0.045 |
| suckers vs. l.crab | 1 | 4.004775 | 0.6863153 | 39.38247 | 0.001 | 0.045 |
| suckers vs. d.crab | 1 | 3.742750 | 0.607901 | 27.90678 | 0.001 | 0.045 |
| suckers vs. m.egg | 1 | 1.134559 | 0.247461 | 5.919026 | 0.001 | 0.045 |
| suckers vs. u.egg | 1 | 3.003539 | 0.4886757 | 17.20271 | 0.001 | 0.045 |

| | | | | | | |
|---|---|---|---|---|---|---|
| suckers vs. r.egg | 1 | 3.223832 | 0.5590918 | 22.82483 | 0.001 | 0.045 |
| u.egg vs. l.crab | 1 | 3.408290 | 0.5753842 | 24.39127 | 0.001 | 0.045 |
| u.egg v d.crab | 1 | 3.287067 | 0.5147332 | 19.093 | 0.001 | 0.045 |
| u.egg v m.egg | 1 | 2.262548 | 0.3536562 | 9.848955 | 0.001 | 0.045 |
| u.egg v r.egg | 1 | 3.113996 | 0.4910765 | 17.36877 | 0.001 | 0.045 |
| m.egg vs. l.crab | 1 | 3.444569 | 0.5496118 | 21.96552 | 0.001 | 0.045 |
| m.egg vs. d.crab | 1 | 3.244994 | 0.4878674 | 17.14714 | 0.001 | 0.045 |
| m.egg vs. r.egg | 1 | 2.703895 | 0.4334167 | 13.76938 | 0.001 | 0.045 |
| r.egg vs. l.crab | 1 | 3.935405 | 0.6726917 | 36.99401 | 0.001 | 0.045 |
| r.egg vs. d.crab | 1 | 3.693634 | 0.5965028 | 26.60997 | 0.001 | 0.045 |
| d.crab vs. l.crab | 1 | 3.693867 | 0.6740108 | 37.21656 | 0.001 | 0.045 |

**Supplemental Table 2 | Structural determination of bacterial metabolites.** $^{1}$H and $^{13}$C NMR in DMSO-$d_6$ data.

| positions | $\delta_C$[a] | type | $\delta_H$[b] | mult (*J* in Hz) | |
|---|---|---|---|---|---|
| 1 | 134.6 | C | | | 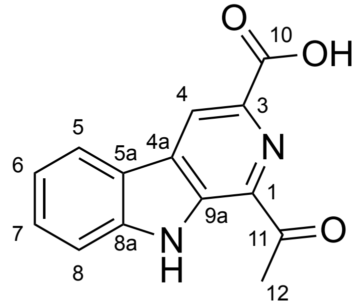 1-acetyl-3-carboxy β-carboline |
| 3 | 134.6 | C | | | |
| 4 | 120.4 | CH | 9.08 | s | |
| 4a | 131.3 | C | | | |
| 5a | 120.4 | C | | | |
| 5 | 122.0 | CH | 8.39 | d (8.0) | |
| 6 | 120.6 | CH | 7.33 | dd (8.0, 8.0) | |
| 7 | 129.0 | CH | 7.60 | dd (8.0, 8.0) | |
| 8 | 113.2 | CH | 7.82 | d (8.0) | |
| 8a | 142.2 | C | | | |
| 9 | | NH | 12.07 | s | |
| 9a | 134.7 | C | | | |
| 10 | 167.2 | C | | | |
| 11 | 201.4 | C | | | |
| 12 | 25.8 | $CH_3$ | 2.84 | s | |
| 1 | 141.4 | C | | | 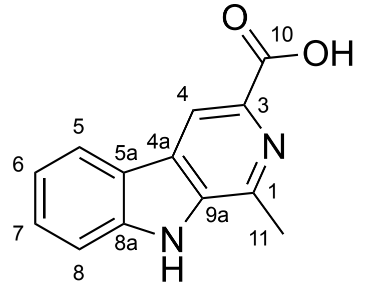 1-methyl-3-carboxy β-carboline |
| 3 | 135.9 | C | | | |
| 4 | 115.1 | CH | 8.73 | s | |
| 4a | 127.1 | C | | | |
| 5a | 121.4 | C | | | |
| 5 | 122.1 | CH | 8.33 | d (8.0) | |
| 6 | 120.0 | CH | 7.29 | dd (8.0, 8.0) | |
| 7 | 128.2 | CH | 7.58 | dd (8.0, 8.0) | |
| 8 | 112.2 | CH | 7.64 | d (8.0) | |
| 8a | 140.8 | C | | | |
| 9a | 135.9 | C | | | |
| 10 | 167.2 | C | | | |
| 12 | 20.2 | $CH_3$ | 2.83 | s | |
| 1 | | NH | 11.67 | d (1.5) | 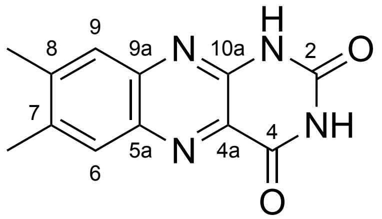 lumichrome |
| 2 | 146.4 | C | | | |
| 3 | | NH | 11.84 | d (1.5) | |
| 4 | 160.6 | C | | | |
| 4a | 130.2 | C | | | |
| 5a | 138.3 | C | | | |
| 6 | 128.6 | CH | 7.93 | s | |
| 7 | 138.8 | C | | | |
| 8 | 144.6 | C | | | |
| 9 | 125.8 | CH | 7.71 | s | |
| 9a | 141.6 | C | | | |
| 10a | 150.1 | C | | | |
| 11 | 20.2 | $CH_3$ | 2.50 | s | |
| 12 | 19.6 | $CH_3$ | 2.46 | s | |

[a]150 MHz and [b]600 MHz, respectively.

**Supplemental Table 3 | Output of Hill equation fits.** Nonlinear fit values were produced in PRISM using the four parameter variable slope model. $Y=Bottom + (X^{Hill\ slope})*(Top-Bottom)*(X^{Hill\ slope}+ EC50^{Hill\ slope})^{-1}$

| | Bottom | Hill Slope | Top | EC50 | $R^2$ | Constrained |
|---|---|---|---|---|---|---|
| nootkatone | 1.663 | 1.829 | 7.255 | 20.33 | 0.8584 | no |
| tryptophan | 1.116 | 2.4 | 1.5 | 554.2 | 0.8151 | yes |
| 1234thh3ca | 1.192 | 1.779 | 2 | 1219 | 0.4056 | yes |
| 1S3S1A3 | 1.331 | 2.107 | 5.5 | 105.8 | 0.9321 | yes |
| 1R3S1A3 | 0.8164 | 1.473 | 5.356 | 83.38 | 0.6605 | no |
| 1234th9hp34bi | 1.357 | 4.811 | 2.422 | 505.1 | 0.3034 | no |
| harmine | 2.042 | 2.603 | ~ 9.100 | 46.33 | 0.5668 | yes |
| harmane | 1.797 | 14.34 | 13.8 | 102.9 | 0.7444 | no |
| norharmane | 1.541 | 2.583 | 9.797 | 77.29 | 0.6612 | no |
| 1A3 | 1.321 | 2.604 | 6.661 | 313.8 | 0.9512 | no |
| H3C | 1.323 | 2.571 | ~ 10.00 | 223.9 | 0.773 | yes |
| Harmol | 1.797 | 14.34 | 15 | 102.9 | 0.7324 | yes |
| 8a-oxo | 3.509 | 1.656 | 29.01 | 171.2 | 0.8503 | no |
| Riboflavin | 1.035 | 2.341 | 2.072 | 57.21 | 0.8442 | no |
| Roseoflavin | 1.405 | 2.914 | 15 | 371.9 | 0.9449 | yes |
| 24da67dipp | 1.67 | 2.28 | 4.8 | 35 | 0.7616 | yes |
| 78dmbgp241h3hd | 1.247 | 11.66 | 2.043 | 265.9 | 0.7442 | no |
| lumichrome | 1.143 | 2.31 | 5.075 | 49.39 | 0.4897 | no |
| 1389tmbgp24d | 1.659 | 2.033 | ~ 16.50 | 16.15 | 0.3166 | yes |
| alloxazine | 1.044 | 1.189 | 2.732 | 637.9 | 0.1952 | no |

**Supplemental Table 4 | Cryo-EM data collection and refinement statistics for CRT1-agonist complexes.**

| | CRT1-Norharmane | CRT1-H3C | CRT1-Lumichrome |
|---|---|---|---|
| **Data collection** | | | |
| EM facility | UCSD | UCSD | UCSD |
| Magnification (kx) | 130 | 81 | 81 |
| Voltage (kV) | 300 | 300 | 300 |
| Electron exposure ($e^-/Å^2$) | 50 | 50 | 50 |
| Defocus range (µm) | -1.4 to -2.4 | -1.4 to -2.2 | -1.4 to -2.2 |
| Pixel size (Å) | 0.935 | 1.072 | 1.072 |
| Micrographs | 3,425 | 8,886 | 14,536 |
| **Reconstruction** | | | |
| Initial particle number | 674,485 | 3,061,405 | 5,833,121 |
| Final particle number | 43,339 | 60,658 | 44,653 |
| Symmetry | C5 | C5 | C5 |
| Box size (pixels) | 320 | 320 | 320 |
| FSC threshold | 0.143 | 0.143 | 0.143 |
| Map resolution (Å) | 3.13 | 3.04 | 3.28 |
| Map sharpening B factor ($Å^2$) | 114.6 | 110.0 | 108.7 |
| **Refinement** | | | |
| Molprobity score | 1.51 | 1.20 | 1.23 |
| Clash score | 4.11 | 2.75 | 3.40 |
| Poor rotamers (%) | 0.00 | 0.00 | 0.00 |
| RMSD values | | | |
| Bond lengths (Å) | 0.005 | 0.006 | 0.006 |
| Bond angles (°) | 0.910 | 0.940 | 0.890 |
| Ramachandran plot | | | |

| | | | |
|---|---|---|---|
| Favored (%) | 95.52 | 97.24 | 97.52 |
| Allowed (%) | 4.48 | 2.76 | 2.48 |
| Outliers (%) | 0.00 | 0.00 | 0.00 |
| B factors (Å$^2$) | | | |
| Protein | 184 | 140 | 175 |
| Ligand | 186 | 148 | 181 |
| Model composition | | | |
| Non-hydrogen atoms | 24,755 | 24,595 | 24,765 |
| Protein residues | 1,460 | 1,460 | 1460 |
| N-glycan | 30 | 20 | 25 |
| Ligand | 5 | 5 | 5 |

**Supplemental Table 5 | Chemical metrics for structure-activity relationships.**

| | PubChem ID | XLogP | LogD[c] | HLB[d] | OEN[e] |
|---|---|---|---|---|---|
| nootkatone | 1268142 | 3.9[a] | 3.81 | 2.38 | |
| tryptophan | 6305 | -1.1[b] | -1.57 | 13.66 | 8.22 |
| 1234thh3ca | 73530 | -0.8[a] | -1.42 | 12.96 | 8.44 |
| 1234th9hp34bi | 107838 | 1.5[a] | -0.26 | 11.45 | 8.31 |
| harmine | 5280953 | 3.6[b] | 1.83 | 7.13 | 10.13 |
| harmane | 5281404 | 3.6[b] | 2.09 | 5.5 | 10.24 |
| norharmane | 64961 | 3.2[b] | 1.93 | 6.11 | 10.26 |
| 1A3 | 5488588 | 2.2[a] | -1.32 | 8.05 | 10.40 |
| H3C | 5406157 | 2.6[a] | -0.79 | 7.64 | 10.41 |
| Harmol | 68094 | 0.7[a] | 2.27 | 6.56 | 11.40 |
| 8a-oxo | 17756791 | 0.3[a] | -0.44 | 9.62 | 9.93 |
| Riboflavin | 493570 | -1.5[b] | -2.41 | 12.98 | 10.00 |
| Roseoflavin | 49867612 | 1.7[b] | -2.61 | 13.35 | 10.06 |
| 24da67dipp | 160723 | 1.5[a] | 2.09 | 6.69 | 10.49 |
| 78dmbgp241h3hd | 95561544 | 0.3[a] | -0.36 | 12.35 | 11.22 |
| lumichrome | 5326566 | 1.1[a] | 1.08 | 8.9 | 11.41 |
| 1389tmbgp24d | 71337284 | 1.5[a] | 2.01 | 8.61 | 11.41 |
| alloxazine | 5372720 | 0.4[a] | 0.14 | 14.55 | 11.45 |

[a]= XLogP-AA
[b]=XLogP
[c]= LogD Chemaxon
[d]= HLB Chemaxon
[e]=Orbital electronegativity of β-Nitrogen for β-carboline molecules and the 3'-Nitrogen for flavin molecules

**Supplemental Table 6 | PERMANOVA analysis on decaying crab microbiome Bray Curtis dissimilarity composition.** Data were computed in the R environment using the command "adonis2". Df: degrees of freedom. Sum Sq: Sum of squares. Pr(>F): P-value associated with the F-statistic. P.adjusted: Bonferroni test adjusted P-value.

| PERMANOVA 9999 permutations. Bray Curtis by decay time point. | | | | | |
|---|---|---|---|---|---|
| | Df | Sum Sq | R2 | F-value | Pr(>F) |
| Groups | 8 | 12.803351 | 0.5776771 | 13.84954 | 0.001 |
| Residuals | 81 | 9.360158 | 0.4223229 | | |
| Totals | 89 | 22.163509 | 1.0000000 | | |

| Pairwise comparisons, multiple comparisons corrected with the Bonferroni test | | | | | | |
|---|---|---|---|---|---|---|
| | Df | Sum Sq | R2 | F-value | Pr(>F) | P.adjusted |
| D0 vs. D1 | 1 | 3.137155 | 0.5803671 | 24.89463 | 0.001 | 0.036 |
| D2 vs. D0 | 1 | 3.579149 | 0.6421793 | 32.30542 | 0.001 | 0.036 |
| D2 vs. D1 | 1 | 0.615750 | 0.209393 | 4.767317 | 0.001 | 0.036 |
| D3 vs. D0 | 1 | 3.807872 | 0.6777237 | 37.8527 | 0.001 | 0.036 |
| D3 vs. D1 | 1 | 1.237144 | 0.3661823 | 10.39933 | 0.001 | 0.036 |
| D3 vs. D2 | 1 | 0.604276 | 0.2308419 | 5.402209 | 0.001 | 0.036 |
| D4 vs. D0 | 1 | 3.829154 | 0.6959905 | 41.20868 | 0.001 | 0.036 |
| D4 vs. D1 | 1 | 1.505551 | 0.4290871 | 13.52845 | 0.001 | 0.036 |
| D4 vs. D2 | 1 | 0.8642183 | 0.3332404 | 8.996236 | 0.001 | 0.036 |
| D4 vs. D3 | 1 | 0.4573388 | 0.2283323 | 5.326101 | 0.001 | 0.036 |
| D5 vs. D0 | 1 | 3.686617 | 0.6461756 | 32.87269 | 0.001 | 0.036 |
| D5 vs. D1 | 1 | 1.711800 | 0.4215145 | 13.11574 | 0.001 | 0.036 |
| D5 vs. D2 | 1 | 0.9397345 | 0.311688 | 8.150931 | 0.001 | 0.036 |
| D5 vs. D3 | 1 | 0.7310218 | 0.2787259 | 6.955837 | 0.001 | 0.036 |
| D5 vs. D4 | 1 | 0.5020864 | 0.2225932 | 5.153902 | 0.001 | 0.036 |
| D6 vs. D0 | 1 | 3.843186 | 0.6928869 | 40.61032 | 0.001 | 0.036 |
| D6 vs. D1 | 1 | 1.871314 | 0.479166 | 16.55996 | 0.001 | 0.036 |
| D6 vs. D2 | 1 | 1.259489 | 0.4171167 | 12.88097 | 0.001 | 0.036 |
| D6 vs. D3 | 1 | 1.041523 | 0.3978315 | 11.89196 | 0.001 | 0.036 |
| D6 vs. D4 | 1 | 0.7031372 | 0.3283469 | 8.799547 | 0.001 | 0.036 |
| D6 vs. D5 | 1 | 0.4317179 | 0.1948082 | 4.354923 | 0.001 | 0.036 |
| D7 vs. D0 | 1 | 3.544279 | 0.5782733 | 24.68168 | 0.001 | 0.036 |
| D7 vs. D1 | 1 | 1.652574 | 0.3617745 | 10.2032 | 0.001 | 0.036 |
| D7 vs. D2 | 1 | 1.175553 | 0.3079841 | 8.010962 | 0.001 | 0.036 |
| D7 vs. D3 | 1 | 1.141374 | 0.3171185 | 8.358893 | 0.001 | 0.036 |
| D7 vs. D4 | 1 | 0.8332493 | 0.2642795 | 6.465813 | 0.001 | 0.036 |
| D7 vs. D5 | 1 | 0.5359922 | 0.1674064 | 3.619191 | 0.001 | 0.036 |
| D7 vs. D6 | 1 | 0.4475713 | 0.1599557 | 3.427442 | 0.001 | 0.036 |
| D8 vs. D0 | 1 | 3.817205 | 0.6462698 | 32.88624 | 0.001 | 0.036 |
| D8 vs. D1 | 1 | 2.063944 | 0.4603054 | 15.35219 | 0.001 | 0.036 |
| D8 vs. D2 | 1 | 1.724282 | 0.4455305 | 14.46346 | 0.001 | 0.036 |
| D8 vs. D3 | 1 | 1.471929 | 0.4285992 | 13.50153 | 0.001 | 0.036 |
| D8 vs. D4 | 1 | 1.149788 | 0.3866174 | 11.34547 | 0.001 | 0.036 |
| D8 vs. D5 | 1 | 0.6940273 | 0.2423027 | 5.756188 | 0.001 | 0.036 |
| D8 vs. D6 | 1 | 0.6080867 | 0.2468755 | 5.900431 | 0.001 | 0.036 |
| D8 vs. D7 | 1 | 0.4398384 | 0.138478 | 2.893256 | 0.004 | 0.144 |
